# Co-option of *Plasmodium falciparum* PP1 for egress from host erythrocytes

**DOI:** 10.1101/2020.01.05.890483

**Authors:** Aditya S. Paul, Alexandra Miliu, Joao A. Paulo, Jonathan M. Goldberg, Arianna M. Bonilla, Laurence Berry, Marie Séveno, Catherine Braun-Breton, Aziz L. Kosber, Brendan Elsworth, Jose S.N. Arriola, Maryse Lebrun, Steven P. Gygi, Mauld H. Lamarque, Manoj T. Duraisingh

## Abstract

Asexual proliferation of the *Plasmodium* parasites that cause malaria follow a developmental program that alternates non-canonical intraerythrocytic replication with dissemination to new host cells. We carried out a functional analysis of the *Plasmodium falciparum* homolog of Protein Phosphatase 1 (*Pf*PP1), a universally conserved cell cycle factor in eukaryotes, to investigate regulation of parasite proliferation. *Pf*PP1 is indeed required for efficient replication, but is absolutely essential for egress of parasites from host red blood cells. A phosphoproteomic screen and chemical-genetic analysis provided evidence for a HECT E3 protein-ubiquitin ligase, as well as a fusion protein with guanylyl cyclase and phospholipid transporter domains, as functional targets of *Pf*PP1. Extracellular phosphatidylcholine stimulates *Pf*PP1-dependent egress. Parasite *Pf*PP1 acts as a master regulator that can integrate multiple cell-intrinsic pathways with external signals to direct parasite egress from host cells.

---

Malaria parasites from the genus *Plasmodium* follow an unusual developmental program during infection of erythrocyte host cells, utilizing a non-canonical style of asexual proliferation, known as schizogony, to undergo multiple cycles of nuclear replication before a single cytokinesis event (*1–3*). Merozoites, the mature parasite forms, must infect new host cells to initiate new intraerythrocytic developmental cycles (IDC) and sustain proliferation, achieved only through egress from the host cell for release into circulation. Protein phosphorylation in parasites is developmentally regulated in blood-stage growth (*4*), and genetic studies show that roughly one-half of the *Plasmodium* protein kinase and protein phosphatase genes are essential for the IDC (*5–16*). Protein Phosphatase 1 (PP1) is a highly conserved and ubiquitous enzyme in eukaryotes that regulates mitotic exit and cytokinesis (*17–21*). With functions also in non-cell cycle-related processes [reviewed in (*22*)], PP1 is a dominant contributor to total cellular phosphatase activity (*23–26*). For the *Plasmodium falciparum* homolog of PP1 (*Pf*PP1) (*27*), genetic evidence for likely essentiality (*6, 8, 28*), high levels of expression compared to other protein phosphatase genes (*29, 30*) (fig. S1A), and the identification of several binding proteins (*31–39*), suggest multiple roles in blood-stage parasites.

To investigate *Pf*PP1-mediated regulation of the IDC and parasite proliferation, we initiated a reverse genetic analysis. With a transgenic *P. falciparum* line expressing a triple-hemagglutinin (HA_3_) tag at the 3’-end of the endogenous *pfpp1* gene (fig. S1B,C), we found that *Pf*PP1 protein is expressed throughout the 48-hr IDC, becoming upregulated from approximately the midpoint until the end (Fig. 1A). In accord with recent reports for PP1 in the rodent malaria parasite *Plasmodium berghei* (*35*), we observed nuclear and cytoplasmic localization of the enzyme, with cytoplasmic *Pf*PP1 becoming predominant as the parasite matures (fig. S1D). Cell cycle stage-dependent compartmentalization of PP1 has been reported in lower eukaryotes, consistent with distinct functions for the enzyme (*40*). To assess the function of *Pf*PP1 in blood-stage proliferation, we generated a transgenic line of *P. falciparum* for inducible knockout of the *pfpp1* gene (*pfpp1-*iKO), based on the dimerizable Cre recombinase (DiCre) system (fig. S1E-G) (*41, 42*). Induction of *pfpp1-*knockout by rapamycin treatment early in the IDC, ∼3-5 hrs post-invasion (hpi), results in strong reduction of *Pf*PP1 protein levels by ∼30 hpi (Fig. 1B; fig. S1B,G). By following DNA replication through the IDC, we found that early-iKO of *pfpp1* delays parasite development before resulting in the accumulation of multinucleate schizont forms, blocked prior to egress (Fig. 1C). Reverse genetic analysis by inducible expression thus establishes the essentiality of *Pf*PP1 for asexual proliferation.

**Fig. 1.**
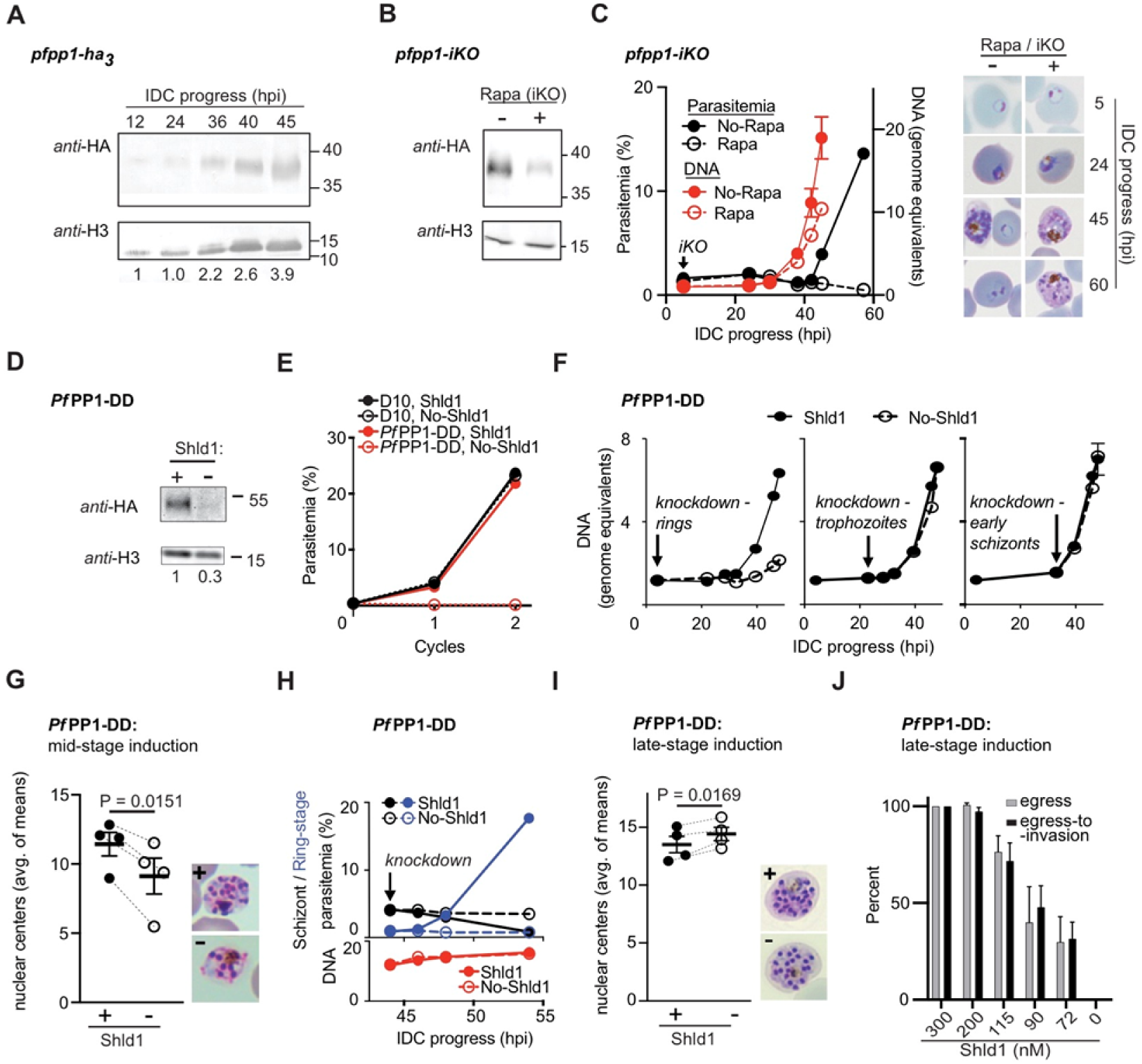
Isolating an essential *Pf*PP1-function late in the intraerythrocytic developmental cycle. **(A)** *Pf*PP1-HA3 expression during intraerythrocytic development (hours post-invasion, hpi), assessed by immunoblot. Relative *Pf*PP1 levels (below lanes) are normalized to histone H3 loading controls. **(B)** iKO of *pfpp1* was initiated at 5 hpi with addition of rapamycin (Rapa) and protein levels were assessed by immunoblot at 30 hpi. **(C)** Left: Parasitemia and DNA synthesis over the IDC following +/-Rapa-treatment at 5 hpi in *pfpp1-iKO* parasites, monitored by flow-cytometry (mean +/− s.d.; n=3 technical replicates). Representative of 4 experiments. Right: Images of parasites along the IDC, following +/-Rapa-treatment at 5-hpi. **(D)** HA-tagged *Pf*PP1-DD protein from schizont-stage parasites (∼48 hpi) grown +/-Shld1 for 6 hrs, assessed by immunoblot. **(E)** Proliferation of *Pf*PP1-DD and parental D10 parasites (wild type), +/-Shld1, monitored by flow-cytometry. Knockdown was induced in cycle-zero. Representative of 2 experiments. **(F)** DNA replication in *Pf*PP1-DD parasites following knockdown (+/-Shld1) at the indicated timepoints and stages of the IDC, monitored by flow-cytometry (mean +/− range; n=2 technical replicates; representative of 3 experiments for early ring induction, 2 experiments for late-ring/trophozoite induction). **(G)** Left: Nuclear centers in terminally developed parasites, assessed by light microscopy, with or without knockdown of *Pf*PP1-DD since the midpoint of the IDC, 22-30 hpi (mean +/− s.e.m.; n=4 experiments). Right: Representative images of terminal parasites +/-Shld1 (Giemsa-stained thin-blood smears). **(H)** Top: Schizont and ring-stage parasites monitored by flow-cytometry following induction of *Pf*PP1-DD knockdown (+/-Shld1) late in the IDC. Bottom: DNA content in the parallel samples, with addition of E64 (50 µM). Representative of 2 biological replicates. **(I)** Nuclear centers following *Pf*PP1-DD knockdown at 44 hpi, as in (G). **(J)** Egress and egress-to-invasion following induction of partial *Pf*PP1-DD knockdown at sublethal doses of Shld1. For this and all remaining panels, egress is based on the rupture of schizonts, and egress-to-invasion is based on the number of ring-stage parasites following experimental treatment late in the IDC.

To investigate *Pf*PP1-function at specific times through the IDC, we used a transgenic *P. falciparum* line for conditional knockdown (fig. S2A-C). Knockdown of *Pf*PP1 fused to a Destabilization Domain (DD)-tag, induced through depletion in culture of the DD-stabilizing small molecule Shield-1 (Shld1) (*11, 43, 44*), confirms essentiality of the enzyme to blood-stage parasites (Fig. 1D,E; fig. S2D,E). We confirmed late IDC-stage expression of the enzyme and cytoplasmic localization in mature parasites (fig. S2F,G). To map the time of function of *Pf*PP1, we induced knockdown at different stages of the IDC and measured DNA replication. We measured phenotypes with knockdown induced at the immature ring stage preceding the growth phase (4 hpi), at trophozoites before the onset of DNA replication (24 hpi), and early in schizogony (33 hpi) (Fig. 1F). With destabilization of *Pf*PP1-DD in rings, we observed significant defects in DNA replication (Fig. 1F; fig. S2H). Knockdown induced in trophozoites or early schizonts permits DNA replication (Fig. 1F; fig. S2I) but results in parasites with reduced numbers of nuclear centers (Fig. 1G; fig. S2J), suggesting defects in nuclear division.

Electron microscopy of *Pf*PP1-DD knockdown parasites indicate failure to complete the terminal mitosis and cytokinesis step of the IDC (fig. S2K), suggesting cell cycle-regulatory functions for *Pf*PP1 conserved from non-parasitic eukaryotes (*19–21*). Defects upstream of cytokinesis with *Pf*PP1-DD knockdown are supported by immunofluorescence analysis; we observe that the inner membrane complex (IMC, antigen *Pf*GAP45) that separates replicated, intracellular parasites fails to form (fig. S2L).

To map functions for *Pf*PP1 late in the IDC, we induced knockdown in schizonts (∼44 hpi), revealing an acute requirement of the phosphatase for egress after complete DNA replication and nuclear segregation (Fig. 1H,I). Knockdown elicits a complete block in host cell egress and erythrocyte re-invasion (Fig. 1H; fig. S2M,N). Partial knockdown late in the IDC elicits sublethal defects in egress without additional defects observed in the further transition to invaded erythrocytes (egress-to-invasion) (Fig. 1J), suggesting a specific and primary function in late stage schizonts.

The effects of late *Pf*PP1-DD knockdown are recapitulated in the *pfpp1-iKO* line with induction of Rapa-mediated iKO later in the IDC (30 hpi, fig. S3A), resulting in depletion of *Pf*PP1 protein at the late schizont stage (Fig. 2A). Late iKO blocks passage to new erythrocytes without defects in DNA replication or nuclear segregation (Fig. 2B,C; fig. S3B). Electron microscopy shows that parasites having undergone late *pfpp1-*iKO display gross morphology typical of maturation, including intact erythrocyte membranes, parasitophorous vacuoles that completely house parasites within, and individual parasite cells physically distinguished by plasma membranes indicating the completion of cytokinesis (Fig. 2D). Immunofluorescence microscopy to image markers for the parasite plasma membrane (antigen *Pf*MSP1) and the underlying IMC (antigen *Pf*MTIP1) confirm cytokinesis and segregation of these structures into replicated parasites (Fig. 2E,F). Immunofluorescence shows also that secretory organelles utilized for invasion, micronemes (antigen *Pf*AMA1) and rhoptry necks (antigen *Pf*RON4), form normally with iKO of *pfpp1* (Fig. 2E,F). In the *Pf*PP1-DD line, immunofluorescence shows that parasites induced for knockdown late in the IDC undergo cytokinesis (IMC antigen *Pf*GAP45) and form normal rhoptries (antigen *Pf*RhopH3) (fig. S3E,F).

**Fig. 2.**
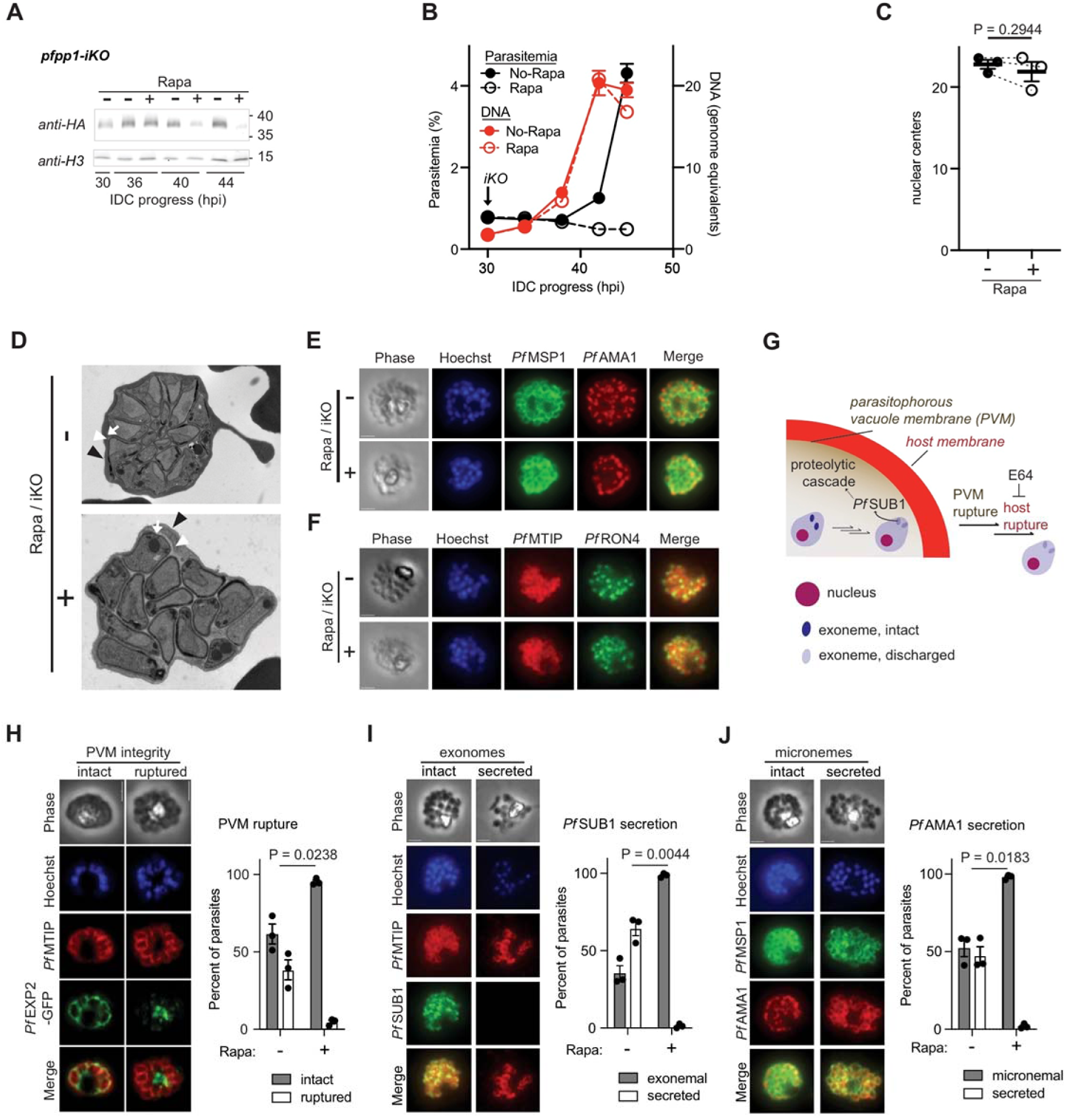
*Pf*PP1 function an early step of parasite egress from erythrocytes. **(A)** *Pf*PP1-HA_3_ expression +/− Rapa-mediated iKO of *pfpp1* at 30 hpi, assessed by immunoblot. **(B)** Parasitemia and DNA synthesis following iKO of *pfpp1* at 30 hpi, as in Fig. 1C (mean +/− s.d.; n=3 technical replicates). Representative of 4 experiments. **(C)** Nuclear centers in terminally developed parasites following Rapa-mediated iKO of *pfpp1* at 30 hpi, as in Fig. 1 (mean +/− s.e.m.; n=3 experiments). **(D)** Electron microscopy of terminally developed *pfpp1*-iKO parasites treated +/− Rapa at 30-hpi. In both images, the different membranes are indicated as follows: erythrocyte (black arrowhead), PV (white arrowhead), and parasite (white arrow). **(E-F)** Immunofluorescence analysis of the microneme antigen *Pf*AMA1 (E) or the rhoptry-neck antigen RON4 (F) in terminally developed parasites +/-iKO of *pfpp1* at 30 hpi. The images also show the parasite plasma membrane marker MSP1 (E) and the inner membrane complex marker MTIP (F). For both apical organelle markers, representative of 2 biological replicates. **(G)** In a mature parasite, regulated secretion of *Pf*SUB1 from exonemes stimulates a proteolytic cascade leading to sequential rupture of the PVM and the erythrocyte host membranes. **(H)** Assessment of PVM rupture at 45 hpi in *pfpp1-iKO* / *Pf*EXP2-GFP parasites treated +/− Rapa at 30 hpi. Left: immunofluorescence images of parasites with intact or ruptured PVMs. Right: Proportion of infected cells exhibiting PVM rupture (mean +/− S.E.M.; n=3 experiments). Parasites in (H)-(J) were treated with E64 (50 µM) at 41 hpi; the completion of cytokinesis was assessed with the inner membrane complex marker *Pf*MTIP or the parasite plasma membrane marker *Pf*MSP1. **(I)** With +/− Rapa-treatment at 30 hpi in *pfpp1-*iKO parasites, quantification of *Pf*SUB1 secretion from exonemal compartments (loss of punctate fluorescence in images at left), as in (H) (mean +/− S.E.M.; n=3 experiments). **(J)** Assessment of AMA1 secretion from micronemes, as in (H-I) (mean +/− S.E.M.; n=3 experiments).

To initiate egress from erythrocytes at the close of the IDC, parasites secrete the protease *Pf*SUB1 in a regulated fashion from exoneme organelles into the lumen of the parasitophorous vacuole and activate a proteolytic cascade for sequential rupture of the vacuolar (PVM) and host cell membranes (*45–47*) (Fig. 2G). To permit assessment of PVM rupture, we endogenously tagged the PVM protein *Pf*EXP2 at the C-terminus with GFP (*48*) in the *pfpp1-iKO* background (fig. S3H-J). Late in the IDC, *Pf*EXP2-GFP in intact PVMs presents intraerythrocytically as a circular label around the parasites or between replicated parasites, while disintegration of the PVM can be observed by the appearance of fluorescent membrane fragments in parasites treated with E64 to prevent host cell rupture (Fig. 2H) (*48*). We used labeling by *Pf*EXP2-GFP to test the requirement for *Pf*PP1 in PVM rupture, finding that late induction of iKO blocks the process in virtually all parasites examined (Fig. 2H). We further used immunofluorescence analysis to assess *Pf*SUB1 release required to initiate PVM rupture (*47*), finding that rapamycin-treated *pfpp1-iKO* parasites fail to secrete the protease from exonemes (Fig. 2I). We observed that discharge of micronemes is also blocked by *pfpp1-*iKO (Fig. 2J). Our findings indicate an essential function for *Pf*PP1 at an early step of egress, following merozoite formation but upstream of discharge of the specialized parasite organelles carrying proteases and other factors utilized for rupture of surrounding membranes.

Our reverse genetic analysis establishing the function of *Pf*PP1 at egress places a factor typically associated with the conventional cell cycle of eukaryotes at a post-replicative stage of the IDC, specifically required for parasitism and spread of infection. To test if dephosphorylation of protein substrates accounts for the requirement for *Pf*PP1 in parasites late in the IDC, we implemented a chemical-genetic approach (*49*) to measure functional interaction between *Pf*PP1 and calyculin A, an active-site inhibitor of eukaryotic PP1 (*50, 51*) (Fig. 3A). We found that knockdown significantly increases the sensitivity of egress-to-invasion by parasites to calyculin A but not to the control antimalarial drug dihydroartemisinin (DHA) (Fig. 3B), supporting a role for *Pf*PP1 phosphatase activity for function. To identify potential substrates of *Pf*PP1 and probe regulation of parasites in the late IDC, we carried out a global phosphoproteomic analysis of *Pf*PP1-DD function spanning a period in the IDC from post-replication through egress, collecting samples from highly synchronized (+/-45 min) *Pf*PP1-DD-intact and knockdown parasites at 48-hpi (4 hrs induction), and at 55-hpi (11 hrs induction) (Fig. 3C). We identified a total of 4720 phosphorylation sites from 1170 phosphoproteins, indicating phosphorylation of ∼1/3 of all *P. falciparum* proteins detected in late-stage parasites (Fig. 3D; Tables S1-S3). We detected a comparable or greater number of phosphorylation sites than totals reported in other phosphoproteomic studies of late-stage *P. falciparum* (*12, 52–55*). Our dataset thus provides a comprehensive view of phosphorylation events with tight time-resolution through the course of egress. At the earliest timepoint following induction of *Pf*PP1-DD knockdown in late in the IDC, there are minimal changes in either the global proteome or phosphoproteome between Shld1-supplemented and knockdown samples, indicating the absence of widespread global changes that may complicate assessment of specific phosphatase functions (Fig. 3D; see Supplementary Text related to figs. S4A-C).

**Fig. 3.**
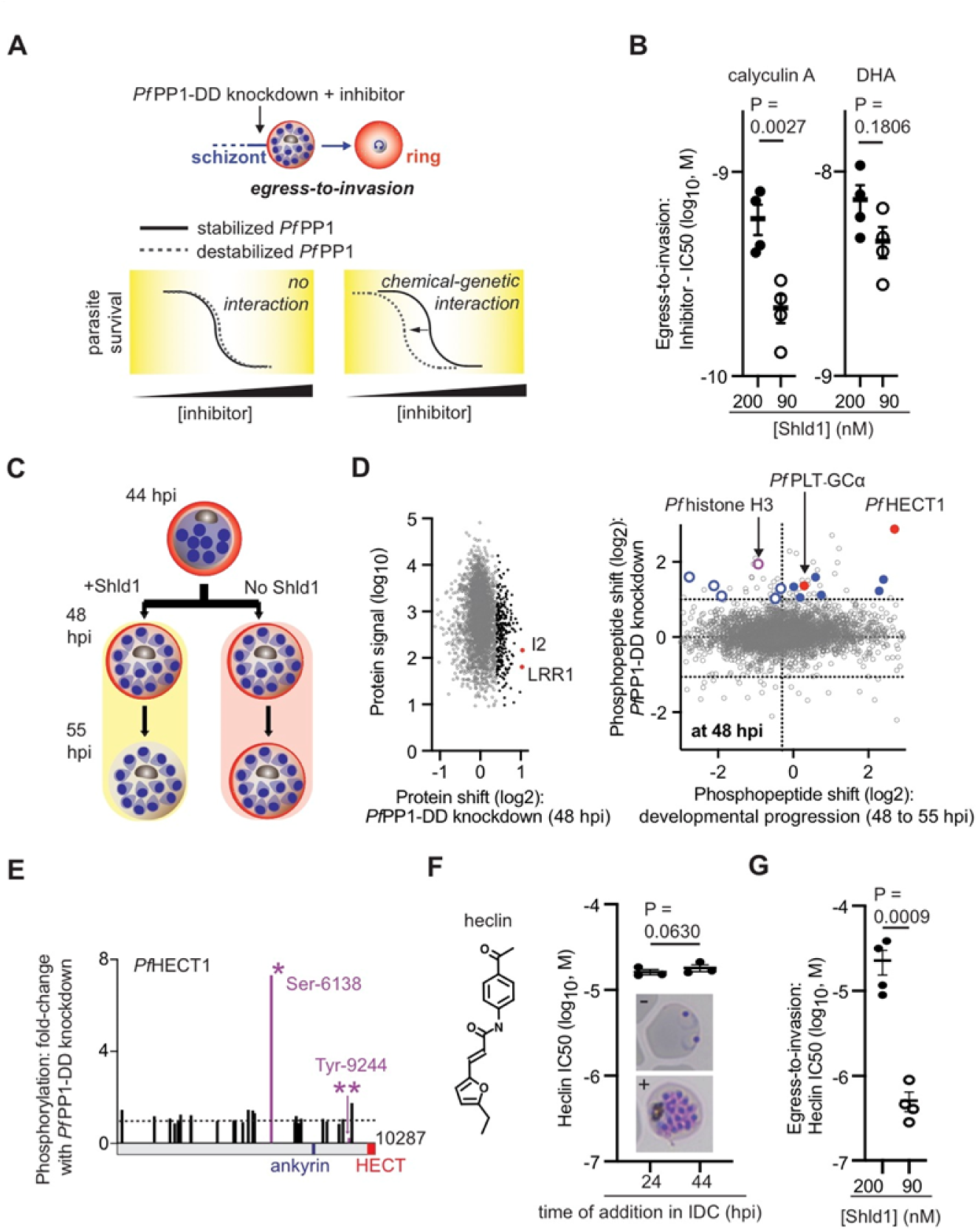
*Pf*PP1 regulation of a HECT E3 protein ubiquitin-ligase for egress. **(A)** Chemical-genetics of *Pf*PP1-DD. We assessed the influence of inhibitors on *Pf*PP1-mediated egress-to-invasion (Fig. 1J) following administration to parasites late in the IDC. To test for functional interactions between *Pf*PP1 and processes targeted by inhibitor, we measured shifts in chemical sensitivity with partial destabilization of the DD-protein. **(B)** The sensitivity (IC50) of *Pf*PP1-mediated egress-to-invasion to the indicated inhibitors at 200 or 90 nM Shld1 (mean +/− s.e.m.; n=4 experiments). **(C)** Scheme for phosphoproteomic analysis of *Pf*PP1-DD knockdown late in the IDC, with samples obtained at 48- and 55-hpi. **(D)** Left: At 48 hpi, levels of individual proteins (summed signal-to-noise) and shift in intensity with *Pf*PP1-knockdown. *Pf*Inhibitor 2 (I2) and *Pf*Leucine rich repeat (LRR) protein 1 are indicated in red, with the top 5% of upregulated proteins indicated in black. Right: For all phosphopeptides detected in late-stage parasites, a plot of changes in levels with *Pf*PP1-DD knockdown at 48-hpi (y-axis) versus changes with development from 48 to 55-hpi in parasites on-Shld1 (x-axis). Two-fold increased phosphorylation (log_2_,1) with knockdown is indicated. Up-phosphorylated phosphopeptides from gene products increased in transcription at the schizont-stage (*30*) are colored; phosphopeptides in the upper-right quadrant least likely to be affected by secondary, developmental-progression defects (see Materials and Methods) are indicated with filled circles. Phosphorylation sites from *Pf*histone H3, *Pf*HECT1, and *Pf*PLT-GCα, are marked in red. **(E)** Schematic of the *Pf*HECT1 protein encoded by PF3D7_0628100, with predicted domains. We show all phosphosites detected in our study with magnitude of change with *Pf*PP1-DD knockdown at 48 hpi (Fig. 3D). The most increased (Ser-6138) and decreased phosphorylation sites (Tyr-9244) are indicated with symbols [*] and [**], respectively. **(F)** Sensitivity of *Pf*PP1-DD parasites on-Shld1 (0.5 µM) to heclin administered at the midpoint (24 hpi) or late in the IDC (44 hpi), determined from erythrocyte re-invasion (mean +/− s.e.m.; n=3 experiments). Representative images of parasites at 55-hpi with or without heclin administration (100 µM) at 44 hpi, are shown below datapoints. **(G)** The sensitivity of *Pf*PP1-mediated egress-to-invasion to heclin, as in (B) (mean +/− s.e.m.; n=4 experiments).

*P. falciparum* homologs of established PP1 regulators for cell cycle progression (*18, 19, 56–59*), the nuclear protein sds22 [in parasites, termed LRR1 for leucine-rich repeat protein 1 (*39*)] and inhibitor-2 (I2) (*38*), are among the two most strongly increased factors in protein expression upon *Pf*PP1-knockdown (Fig. 3D). Perturbed expression of these regulators may indicate a conserved function mediated by parasite *Pf*PP1, while an increase in factors for glycolysis and the pentose phosphate pathway suggest association with the proliferative state of parasites (Table S4) (*60*). At 48 hpi, we observe by both phosphoproteomics and separately by immunoblot analysis increased phosphorylation (by up to ∼5-fold) of Ser-29 of *Pf*histone H3 (Fig. 3D; fig. S4D), homologous to Ser-28 in the human ortholog studied as a phosphorylation site targeted by PP1 during mitotic exit and interphase (*61, 62*). We thus observe among both putative regulators and substrates evidence for conserved PP1 activity in *P. falciparum*.

The 50 proteins increased >2-fold in phosphorylation upon *Pf*PP1-DD knockdown (Fig. 3D, Table S3) include chromatin factors (histone H3 variant, histone deacetylase 1, chromodomain-binding protein), transcription factors from the AP2 family, and vacuolar-protein-sorting family proteins (VPS11 and VPS18). To focus on potential substrates for essential *Pf*PP1 function at egress, we identified gene products that specifically increase in transcriptional expression late in the IDC (Fig. 3D). The top hit from our phosphoproteomic screen, based on magnitude of change in phosphorylation with *Pf*PP1-knockdown, is a previously uncharacterized, late IDC-stage protein carrying a ∼300-amino acid HECT E3 protein-ubiquitin ligase domain at the C-terminus (PF3D7_0628100) (Fig. 3D,E; fig. S4E). In addition to the highly upregulated phosphorylation site (Ser-6138), knockdown of *Pf*PP1-DD also reveals a strongly downregulated site (>4-fold, Tyr-9244); the protein also contains a previously predicted site for interaction with *Pf*PP1 (*34*) near the HECT domain (Fig. 3E; fig. S4E). At >10,000 amino acids, the protein is the largest in the *P. falciparum* proteome (Fig. 3E; fig. S4E).

The large protein, which we name here *Pf*HECT1, is the one HECT domain-containing gene of 4 total in *P. falciparum* to become increased in expression late in the IDC (fig. S4E). To test for specific HECT activity in parasites late in the IDC, we tested for susceptibility to the small molecule heclin (Fig. 3F). Heclin was identified as a broad-spectrum inhibitor of mammalian HECT enzymes, with biophysical studies suggesting that direct binding by the compound interferes with conformational changes necessary to catalyze transfer of ubiquitin from the E2 adaptor protein to an active-site cysteine in the E3 ligase (*63*). In *Pf*PP1-DD parasites stabilized with Shld1, we established the anti-malarial activity of heclin toward parasites, with short (∼4 hrs) and longer periods of exposure (∼24 hrs) preceding egress exhibiting similar potency toward establishment of ring-stage parasites (IC50 ∼20 µM), indicating major inhibitory activity in schizonts (Fig. 3F). While *Pf*PP1-DD destabilization does not increase DNA replication defects induced by heclin (fig. S4F), knockdown in parasites late in the IDC strongly increases susceptibility of egress-to-invasion to the inhibitor, reducing the IC50 by >100-fold to submicromolar levels (Fig. 3G; fig. S4G). An assessment of schizont rupture confirms specific inhibition by heclin toward egress, with dependence on *Pf*PP1 (fig. S4H). Phosphoproteomic analysis (Fig. 3D) with chemical-genetic analysis (Fig. 3G) indicates a critical role for *Pf*PP1 in activation of *Pf*HECT1-mediated E3 protein-ubiquitin ligase activity.

Among other late IDC-stage *P. falciparum* genes showing increased phosphorylation with *Pf*PP1-knockdown is a guanylyl cyclase (GC) domain-containing protein encoded by the gene PF3D7_0381400 (Fig. 3D, 4A), required for production of cyclic guanosine monophosphate (cGMP) to stimulate the downstream effector *Pf*Protein Kinase G (*Pf*PKG) and egress from infected erythrocytes (*9, 47, 64*). Termed GCα, the protein is present across apicomplexan parasites, and in *Toxoplasma gondii* is also essential for egress (*65–68*). Given the function we identified for *Pf*PP1 in egress, we implemented chemical-genetics to test for functional interaction with components of cGMP-mediated signal transduction. To artificially raise cellular cGMP in parasites late in the IDC, we used zaprinast, a cGMP-specific phosphodiesterase (PDE) inhibitor (*47, 69, 70*) (Fig. 4A). The susceptibility of egress-to-invasion by *Pf*PP1-DD parasites to zaprinast becomes sharply increased with knockdown: IC50 values drop by as much as ∼400-fold to submicromolar levels (Fig. 4A; fig. S5B). Functional interaction of *Pf*PP1 with the *Pf*PKG inhibitor Compound-1 (Cpd1) (*9*), by contrast, is weak, with measurements of egress-to-invasion showing no chemical-genetic interaction (Fig. 4A). An assessment of egress shows increased sensitivity to Cpd1 with *Pf*PP1-DD knockdown, though much less than the level of functional interaction observed with zaprinast (Fig. S5C). Our analysis of parasites late in the IDC indicates a primary function for *Pf*PP1 in suppression of guanylyl cyclase activity, restricting cGMP-mediated activation of *Pf*PKG upstream of numerous egress and invasion processes (*54, 71*). We note that in parental, genetically unmodified parasites, Shld1 does not affect sensitivity to calyculin A, heclin, or zaprinast (fig. S5D), indicating that the chemical-genetic interactions we have observed here with *Pf*PP1-DD knockdown result from impairment of phosphatase function.

**Fig. 4:**
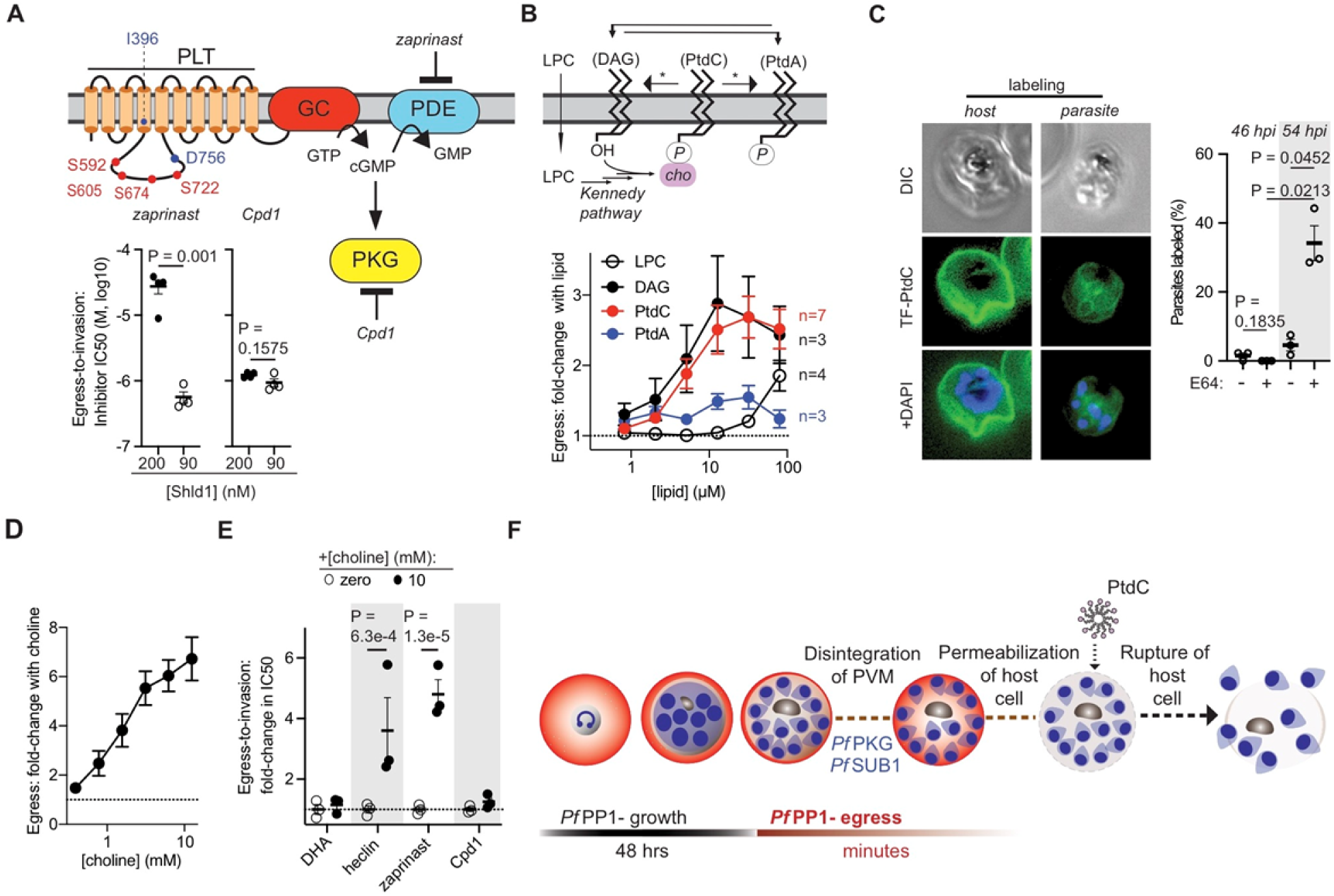
Regulation of egress by extracellular phosphatidylcholine. **(A)** Top: *Pf*PP1-regulated phosphorylation and cGMP-based signal transduction initiated by the parasite fusion protein carrying a phospholipid transporter (PLT) and a guanylyl cyclase (GC) domain (*Pf*PLT-GCα, PF3D7_1138400). We indicate *Pf*PP1-regulated sites in the PLT domain (red) and signature sequences for phospholipid translocase activity (purple), Ile-396 and Asp-756. cGMP-phosphodiesterase (PDE) and *Pf*PKG, as well as inhibitory small molecules for both targets, are depicted. Bottom: The sensitivity of *Pf*PP1-mediated egress-to-invasion to zaprinast (left) or Cpd1 (right), as in Fig. 3 (mean +/− s.e.m.; n=4 experiments). **(B)** Top: Phospholipids at the parasite plasma membrane. Interconversion of DAG and PtdA are likely mediated by parasite homologs of DAG-kinase and phosphatidic acid phosphatase (*73, 74*). Phospholipase enzymes for conversion of PtdC to DAG and PtdC to PtdA (indicated with asterisks) have not been identified in *Plasmodium falciparum* (*88*); PtdC levels are correlated with PtdA (*78*), indicating linkage between the species. Extracellular LPC crosses the plasma membrane, providing substrate for endogenous biosynthesis of PtdC via the Kennedy Pathway (*78, 79*). Bottom: Stimulation of egress by exogenously administered lipids in parasites late in the IDC with partial destabilization of *Pf*PP1-DD (100 nM Shld1), expressed in terms of fold-change relative to egress without lipid (mean +/− s.e.m., with number of separate experiments indicated in plot). **(C)** Labeling of the parasites by PtdC. Left: Representative images of late-stage, *P. falciparum-* infected erythrocytes (*Pf*PP1-DD with 0.5 µM Shld1 and 50 µM E64, 54 hpi) labeled by TF-PtdC at the erythrocyte membrane (host) or at the plasma membranes of internal parasites. Nuclei are indicated by Hoechst dye. Right: The proportion of infected erythrocytes with labeled parasites at the indicated timepoints, +/-E64 (mean +/− s.e.m.; n=3 experiments). **(D)** Stimulation of *Pf*PP1-mediated egress with supplementation of choline, as in (B). **(E)** Modulation of *Pf*PP1-regulated cellular processes by supplemented choline. For each inhibitor, fold-change in IC50 with additional choline in *Pf*PP1-DD parasites in 150 nM Shld1 is indicated (mean +/− s.e.m.; n=3 experiments; multiple *t-*tests). **(F)** A model for the function of *Pf*PP1 through the IDC. Following activity in growth during the IDC, *Pf*PP1 is essential for the egress program upstream of *Pf*PKG and the *Pf*SUB1-initiated proteolytic cascade required for disintegration of the PVM. PtdC enters host cells following natural permeabilization of the erythrocyte membrane, acting on exposed parasites to stimulate egress.

The protein GCα implicated by our analysis is fused at its N-terminus to a putative P4-ATPase phospholipid transporter (PLT) domain predicted to translocate phospholipids from the exoplasmic to the cytoplasmic face of membranes. The fusion of PLT with the GC domain is a structure found only in alveolates (*72*), and we observed that *Pf*PP1-responsive phosphorylation sites (1.4-2.6-fold upregulation with knockdown) cluster in a parasite-specific cytoplasmic loop of PLT containing a catalytic site predicted essential for P4-ATPase activity (Fig. 4A; fig. S5A). Mutational analysis in *T. gondii* GCα indicates the requirement for fusion of the PLT and GC domains as well as ATP-dependent catalysis by the PLT (*65, 66*), raising the possibility of a role for phospholipids in proper function of the protein, perhaps in egress. To directly assess involvement of phospholipids in *Pf*PP1-regulated egress, we administered synthetic phospholipids to parasites late in the IDC. We tested phosphatidic acid (PtdA), known to stimulate host cell egress in *T. gondii* when added extracellularly (*65*) and also through endogenously synthesized forms resulting from intracellular phosphorylation of the neutral lipid diacylglycerol (DAG) (*73, 74*). We also tested phosphatidylcholine (PtdC), the major species of phospholipid in host serum at concentrations of ∼1-2 mM (*75, 76*). In parasites with intact *Pf*PP1-DD (300 nM Shld1), neither phospholipid influences egress (fig. S5E). In parasites with partially destabilized *Pf*PP1-DD, however, PtdA and PtdC stimulate egress by up to ∼1.5 and ∼3-fold, respectively, with effects observed at as low as 5 µM phospholipid (Fig. 4B). An effect for PtdA might be explained by the proposal that the molecule provides one of multiple signals required for efficient egress, acting on targets to activate microneme secretion and parasite motility, downstream of cGMP and phosphatidylinositol signaling (*71, 74, 77*). We indeed observed that knockdown of *Pf*PP1-DD late in the IDC increases susceptibility to the DAG-kinase inhibitor R59022 that restricts endogenous PtdA synthesis (fig. S5F), consistent with a pro-egress function for the phospholipid in *Plasmodium* parasites (*73*).

A role for PtdC in egress, has not been described. We found that extracellular DAG stimulates egress to similar levels at similar doses as PtdC, suggesting convergent targets or efficient conversion between the two lipids upon incorporation into the parasite from the extracellular medium (Fig. 4B). In contrast, lysophosphatidylcholine (LPC), which drives parasite PtdC biosynthesis via the Kennedy Pathway (*78, 79*), stimulates egress weakly in comparison to direct administration of the phospholipid (Fig. 4B); and glucose supplementation to increase endogenous DAG (*78, 80*) does not stimulate egress (Fig. S5G).

While erythrocyte membranes housing developing *P. falciparum* parasites block access to free PtdC (*78*), host cells abruptly become permeable to extracellular solutes in the seconds to minutes preceding egress (*46, 48, 81*), presenting a route for direct interaction between parasites and circulating phospholipids. We thus tested accessibility of parasites to PtdC during egress, using a fluorescent analog (labeled with TopFluor, TF). In most conditions we observed that TF-PtdC marks only the outer membranes of erythrocytes housing multinucleate schizonts (Fig. 4C). We observed clear labeling of parasites within infected erythrocytes, however, only at a later timepoint when parasites have naturally entered the egress program and acquired host cell permeability (Fig. 4C). The intact schizont with permeable erythrocyte membrane is a transient state prior to egress, stabilized by E64-treatment (*46, 81–83*). We conclude that circulating PtdC accesses parasites when host cells become permeable to the extracellular environment, shortly before natural host cell rupture.

Endogenous PtdC is promoted with the addition of the precursor choline (*78, 79*). We found that choline stimulates egress in parasites with partially destabilized *Pf*PP1-DD; albeit in contrast to PtdC, only at non-physiological serum levels (*84*) (Fig. 4D). Identical, high concentrations of choline do not influence egress by parasites with intact *Pf*PP1-DD (300 nM Shld1) (fig. S5H). We found that high choline decreases susceptibility of parasites specifically to heclin and zaprinast (Fig. 4E). Our analysis suggests that the extrinsic PtdC signal for egress interacts with cell-intrinsic pathways regulated by *Pf*PP1.

We have functionally characterized *Pf*PP1 through the blood-stage IDC of malaria parasites. In addition to potential conserved roles in development, *Pf*PP1 is absolutely essential for egress. At the pre-erythrocytic liver-stage, analysis by other researchers shows the non-essentiality of PP1 for intrahepatocytic development, though a function for the phosphatase in egress into the bloodstream was not assessed (*28*). *Pf*PP1 regulates egress of parasites through multiple pathways including *Pf*HECT1 (fig. S6). Chemical inhibition by heclin (*63*) suggests a role for ubiquitination alongside well-established pathways for phosphorylation and proteolysis. At multiple stages of the *Plasmodium* lifecycle, signaling by cGMP is utilized for colonization of new host niches, and studies indicate that a specific timing of activation and level of the second messenger are critical for infectivity (*47, 85–87*). The use of *Pf*PP1 at the blood-stage to suppress cGMP upstream of *Pf*PKG and *Pf*SUB1 may suggest an interface between the parasite maturation and egress programs (Fig. 4F; fig. S6). We additionally discovered extracellular PtdC as an extrinsic serum factor to stimulate egress from erythrocytes, distinct from the extracellular LPC signal that suppresses differentiation to sexual-stage forms at an earlier point in the IDC (*78*). We propose that *P. falciparum* uses PLT-GCα to translocate PtdC from the serum across the parasite plasma membrane to provide a late signal to stimulate egress (Fig. 4F; fig. S6). Our study demonstrates that the *Plasmodium* homolog of PP1 is a regulatory nexus for egress from erythrocytes, balancing cell-intrinsic pathways with environmental signals to ensure release of invasive parasites into circulation and infection of new host cells.

## Methods

### Reagents and antibodies

Rapamycin (LC laboratories), E64 (Sigma-Aldrich, Cat. No E3132); dihydroartemisinin (Sigma-Aldrich, Cat. No. D7439), calyculin A (Sigma-Aldrich, Cat. No. C5552), heclin (Sigma-Aldrich, Cat. No. SML1396), zaprinast (MP Biomedicals, Cat. No. ICN15693180), were each prepared in DMSO. Choline chloride (Sigma-Aldrich, Cat. No. 7527) was prepared in water. DAG and all phospholipids with mono-unsaturated diacylglycerol backbone (16:0, 18:1) from Avanti [PtdA, Cat. No. 840857; PtdC, Cat. No. 850457; TopFluor-PtdC, Cat. No. 810281; and DAG, Cat. No. 800815] were solubilized at 1 or 5 mM in 100% methanol, except PtdC (water: ethanol: methanol; 1:1:2). LPC (Cat. No. 855675) was solubilized at 200 mM in a 1:1 mixture of ethanol and water. Compound-1 (DMSO-based) was a gift from Dr. Jeffrey Dvorin (Boston Children’s Hospital). Dilutions and sources for antibodies for immunoblot or immunofluorescence analysis are as follows: rabbit anti-GAP45 (1:5000, gift from Dr. Julian Rayner, Wellcome Trust Sanger Institute, Hinxton, UK); mouse anti-RhopH3 (1:200, gift from Jean-Francois Dubremetz); rabbit α-MTIP (1:500, gift from Tony Holder, The Francis Crick Institute, UK), mouse α-MSP1.19 (1:1000, gift from M. Blackman, The Francis Crick Institute, UK), mouse α-RON4 [1:200, home-made (*89*)], mouse α-SUB1 (1:2, gift from M. Blackman), rabbit α-AMA1 (1:1000, gift from M. Blackman); rabbit anti-histone H3 (1:10,000, Abcam); rat anti-HA antibody 3F10 (1:1000, Roche); rabbit anti-*Pf*aldolase-HRP (Abcam, 1:2000), mouse anti-GFP (Roche, 1:1000). Shield-1 (Shld1) was synthesized as described (*43, 90, 91*) and dissolved to 1 mM stock concentration in absolute ethanol before use.

### Plasmids

Primers for PCR amplification and verification of transgenesis in *P. falciparum* are shown in Table S5. Transgenic parasites *pfpp1*-HA_3_ were obtained using the plasmid pLN-PP1-HA_3_-loxP. To generate the plasmid pLN-PP1-HA_3_-loxP, we first introduced between the BamHI and HpaI sites in pLN-ENR-GFP (*92*) a synthetic fragment with sequence for a triple hemagglutinin tag (HA_3_) followed by a stop codon and a loxP site (IDT DNA), and multiple cloning sites upstream of the tag. The resulting plasmid pLN-PP1-HA_3_-loxP was further modified to target endogenous *pfpp1* with HA_3_ and loxP by introduction of a 5’ homology region for the gene (HR1, 682 bp of genomic DNA sequence for exons 2 and 3) fused to a recodonized synthetic fragment (IDT DNA) for exons 4 and 5. The PCR-amplified elements were ligated in a single reaction step by In-Fusion cloning (Clontech) upstream of the HA_3_ tag in XmaI and AfeI sites of pLN-HA_3_-loxP. The *pfpp1* 3’ homology region (HR2) carrying 440 bp of the *pfpp1* 3’-UTR was PCR-amplified and inserted by In-Fusion reaction 3’ of the loxP site between PstI and HpaI. The guide RNA sequence for targeting *pfpp1* near exon 3 was cloned into the BbsI sites in pDC2-cam-co-Cas9-U62-hDHFR (gift from M. Lee). For subsequent engineering of a *pfpp1* conditional knockout in parasites, the pLN-PP1-loxPint plasmid was modified with introduction of the following elements ligated in 5’ to 3’ order between the BamHI and ApaI sites of pLN-ENR-GFP: PCR-amplified fragment of *pfpp1* encompassing 5’UTR and part of exon 1 (382 bp), a synthetic fragment for a recodonized 3’ sequence of exon 1 followed by the artificial loxPint (IDT DNA), and a PCR-amplified fragment of the 5’ end of *pfpp1* exon 2 (601 bp). The plasmid for the guide RNA targeting *pfpp1* near exon 1 was constructed as above. To endogenously tag *Pf*EXP2 with GFP, we generated plasmid pLN-*Pf*EXP2-GFP as described (*48*). Two homology regions for the gene *pfexp2* were cloned in pLN-ENR-GFP on both sides of the GFP coding sequence: 549 bp of *pfexp2* 3’ coding sequence without the stop codon in frame with GFP, and 453 bp of *pfexp2* 3’UTR downstream of GFP. The *pfexp2* guide RNA sequence (*48*) was cloned into the BbsI sites of pDC2-cam-co-Cas9-U62-hDHFR. Plasmid pAK8 for 3’-single crossover HA-DD-tagging at endogenous *pfpp1* was constructed in the pJDD41 background (*11*) with the PCR-amplified genomic insert ligated between the NotI and XhoI restriction sites. All plasmid sequences were verified before downstream applications.

### Parasite culture, transfection, and synchronization

D10 or 3D7 (Walter and Eliza Hall Institute), or p230p-based parasites (*41*) were cultured continuously in human erythrocytes (*93*) obtained from a commercial source (Research Blood, Boston) or anonymous donors from the French Bloodbank (Etablissement Français du Sang, Pyrénées Méditerranée, France). Continuous culture was typically carried at 2-5%-hematocrit in RPMI-1640 (Sigma Aldrich, Cat. No. R6504) supplemented with HEPES, 25 mM; Albumax II, 4.31 mg/ml (Thermo Fisher Scientific), or 10% human serum; sodium bicarbonate, 2.42 mM; and gentamycin (20-25 µg/ml). Parasites were cultured at 37° C in hypoxic conditions (1-5% O_2_, 5% CO_2_, with N_2_ as balance) in modular incubator chambers. Parasites were transfected by electroporation (*94*), and treated with WR99210 (2.5-5 nM) or blasticidin (2.5 µg/ml). *Pf*PP1-DD transfectants were further selected for single-crossover integrants by cycles of on-drug and off-drug treatment (*10, 95*). All transgenic lines were cloned by limiting dilution and genotyped by PCR. Unless otherwise noted, all experiments with *Pf*PP1-DD parasites indicate a line constructed in the D10 background.

We synchronized parasites with heparin (100 units/ml) to define restricted periods of invasion (*96*). Alternatively, we isolated schizonts by magnetic-affinity purification (MACs LS column, Miltenyi, fitted with a 23-gauge needle) or 70% Percoll cushion, and allowed invasion into uninfected erythrocytes for a defined period. Following invasion, we either added heparin to block further invasion, selectively lysed remaining schizonts by sorbitol treatment (5% w/v in double-distilled water), or separated recently invaded rings from unruptured schizonts by magnetic affinity purification (MACs LS with 27-gauge needle).

### Statistical significance testing

We carried out all tests for statistical significance in Prism software (GraphPad). Unless otherwise stated, *P-*values indicate the results of paired, two-tailed *t-*tests.

### Induction of *Pf*PP1-phenotypes through conditional expression

For assays, we induced parasites for either knockout of *pfpp1* [mock-(DMSO) versus Rapa, 10 nM] or knockdown of *Pf*PP1-DD [Shld1 (0.2-0.5 µM) versus ethanol vehicle]. In knockout parasites, we washed away Rapa 4 hrs following addition.

### Immunoblot analysis

For immunoblot analysis, we released parasites from erythrocytes with cold PBS containing 0.1-0.2% saponin, and boiled in SDS-PAGE sample buffer as previously described (*10*). Following electrophoretic separation, proteins were transferred to a nitrocellulose membrane, and immunoblot analysis was carried out using the LI-COR system (Lincoln, USA), or the Chemidoc system (Bio-Rad).

### Microscopy

*Light microscopy.* For quantitative assessment of nuclear centers in terminally developed *Pf*PP1-DD parasites [∼54-60 hpi; +E64 (50 µM) since ∼45 hpi], we collected ∼500,000 infected cells onto glass slides by cytospin centrifugation, followed by fixation in methanol and staining with May-Grünwald-Giemsa. All infected cells encountered in visual fields by conventional light microscopy were counted, typically >80 per sample. In the *pfpp1-iKO* line, we used immunofluorescence microscopy (see below) to identify segmented parasites that stained positive for the antigen *Pf*MTIP before counting nuclei stained with DAPI. Immunofluorescence analysis was carried out essentially as described (*97*). Thin smears were fixed in 4%-paraformaldehyde in PBS for 10 minutes at room-temperature or overnight at 4° C in a humidified chamber followed by extensive washing in buffer; permeabilized with 0.1%-Triton-X-100/PBS for 10 minutes at room temperature before further washing; blocked with 3% BSA/PBS for >1 hr at room temperature or overnight at 4° C; treated with primary antibody overnight at 4° C; washed extensively before treatment with the appropriate Alexa-488 secondary antibody (1:1000 dilution) for 1 hr at room temperature; washed and prepared in DAPI-containing mounting solution for imaging. Images were taken with a Zeiss Axioimager Z2 or AxioObserver.Z1, and processed with Zen blue edition software (Zeiss) or Fiji (*98*). For immunofluorescence assays for PVM rupture or secretion of exoneme or microneme antigens, parasites were treated at 41 hpi with 50 µM E64 and smeared 4-5 hrs later for analysis.

For assessment of TF-PtdC labeling of *P. falciparum*, we treated synchronous (+/-1 hr) *Pf*PP1-DD parasites (+0.5 µM Shld1) with or without E64 (50 µM) for ∼7 hrs before image acquisition at the indicated timepoints. After evaporation of TF-PtdC on the surface of multiplate wells, we added parasites in standard media (2%-hematocrit) for a final concentration of fluorescent label of 100 µM. After ∼30 minutes at 37° C in standard culture conditions, cells were collected and stored at 4°C until imaging carried out over the course of the next ∼2 hrs. Just before imaging, parasites were spotted and mixed with Hoechst dye on coverslips (No. 1.5) pre-treated with Concanavalin-A (Sigma-Aldrich, Cat. No. C5275, 0.5 mg/ml in PBS; spread and dried at 37° C for ∼30 minutes) immediately before sealing by surface tension and dispersion of cell suspension with a glass slide. Images were acquired with a 63x-objective on the Zeiss Axioimager Z2 in the DIC, DAPI, and GFP channels. We scored 40-66 multinucleate, infected cells per condition to estimate internal labeling of parasites.

#### Transmission electron microscopy

For *pfpp1-iKO* cells, we directly added 25% glutaraldehyde (EM grade) to the culture medium to obtain a final concentration of 2.5%. After 10 min incubation at room temperature, we centrifuged the cells and resuspended the pellet in 20 volumes of cacodylate buffer (0.1 M) containing 2.5% glutaraldehyde and 5 mM CaCl_2_.The suspension was left 2 hours at RT before long-term storage at 4°C in fixative until further processing. All the following incubation steps were performed in suspension, followed by centrifugation using a benchtop microcentrifuge. Cells were washed with cacodylate buffer and post-fixed with 1% O_s_O_4_ and 1.5% potassium ferrocyanide in cacodylate buffer for 1 hr. After washing with distilled water, samples were incubated overnight in 2% uranyl acetate in water and dehydrated in graded series of acetonitrile. Impregnation in Epon 812 was performed in suspension on a rotary shaker for 1hr in Epon: acetonitrile (1:1) and 2x 1hr in 100% Epon. After the last step, cells were pelleted in fresh epon and polymerized for 48 hrs at 60° C. 70 nm sections were made with an ultramicrotome Leica UC7, contrasted with uranyl acetate and lead citrate and imaged for transmission electron microscopy (TEM) on a JEOL 1200 EX. All chemicals were purchased from Electron Microscopy Sciences (USA).

TEM analysis of *Pf*PP1-DD parasites was carried out similarly, with some modifications. For fixation, 1 volume of suspended culture (> ∼5 µl packed cell volume) was supplemented with 1 volume of a 2x fixative solution (5% glutaraldehyde, 2.5% paraformaldehyde, 0.06% picric acid in 0.2M cacodylate buffer, pH 7.4), spun briefly at 500g, and stored at 4° C before further processing. Following fixation, cells were washed in water, then maleate buffer before incubation in 2% uranyl acetate (1 hr). Following washes in water, dehydration was done in grades of alcohol (10 min each at 50%, 70%, 90%, and 2x 100%). The samples were then put in propyleneoxide for 1 hr and infiltrated overnight in a 1:1 mixture of propyleneoxide and TAAB Epon (Marivac Canada Inc. St. Laurent, Canada). The following day, the samples were embedded in TAAB Epon and polymerized at 60° C for 48 hrs. Ultrathin sections (about 60nm) were cut on a Reichert Ultracut-S microtome, picked up on to copper grids stained with lead citrate and examined in a JEOL 1200EX or a TecnaiG² Spirit BioTWIN. Images were recorded with an AMT 2k CCD camera.

### Flow-cytometry and analysis

All measurements of parasitemia through flow-cytometry were carried out with staining of fixed cells with SYBR-Green I (Invitrogen) to distinguish DNA-containing parasitized erythrocytes from uninfected, enucleate erythrocytes (*10, 13, 99, 100*). Fixation of *Pf*PP1-DD cell suspension was carried out by addition of >3 volumes of PBS supplemented with paraformaldehyde (4% final concentration) and glutaraldehyde (0.0075-0.015% final concentration), followed by storage at 4° C for >12 hrs before further washes in buffer. For quantitative measurements of cellular DNA, fixed cells were permeabilized with Triton-X-100 (0.1%) and RNase-treated (∼0.3 mg/ml) before staining (*13*). For *pfpp1-iKO* parasites, cells were fixed by addition of an equal volume of PBS-paraformaldehyde (8%); fluorescence was measured with a BD FACS Canto I cytometer. For *Pf*PP1-DD parasites, flow-cytometry was carried out with a MacsQUANT (Miltenyi) on the FITC channel. All flow-cytometry data were analyzed with FlowJo software.

To calculate cellular DNA content, we used fluorescence measurements from uninfected erythrocytes (zero genomes), singly-infected rings (1 genome), and doubly-infected rings (2 genomes), to build standard curves for translation of total fluorescence of an infected erythrocyte population into “genome equivalents.” In some measurements, we applied a background correction across all samples to subtract the contribution of parasite cells that did not advance into DNA replication by 48-hpi (i.e. in +Shld1-conditions). To calculate egress from parasites late in the IDC in an experiment, we used the remaining schizont populations measured at the end of an assay. We treated schizont levels in No-Shld1 conditions (or 50 nM Shld1, fig. S5E,H) as a measure of no-egress (zero), and levels with high-Shld1 as full egress (100%). We similarly calculated egress-to-invasion from parasites late in the IDC using ring-stage parasite levels. Fold-change in egress, as reported in Fig. 4, is the quotient of schizont levels at the end of the assay in the absence of chemical (numerator) to schizonts levels left in the presence of chemical (denominator).

### Proteomic and phosphoproteomic profiling

#### Parasite collection and processing for downstream analysis

Proteomic and phosphoproteomic analysis was carried out as described (*13, 101*), with modifications. *Pf*PP1-DD parasites were synchronized as described above using MACs purification of schizonts and was treated with sorbitol (5% w/v in water) after 1.5 hours of invasion into uninfected erythrocytes to eliminate remaining schizonts and isolate freshly re-invaded rings. Approximately 10×10^10^ synchronous ring-stage *Pf*PP1-HA-DD parasites in Shld1 (0.3 µM) were cultured to 44 hpi with a preceding change of warm media at ∼24 hpi, washed extensively and replated in warm (37° C) complete-RPMI at ∼5×10^6^ parasitized erythrocytes per ml of culture +/-0.3 µM Shld1. At 48 hpi, for each Shld1 condition, 2 technical replicates each with or without Shld1 (∼1×10^10^ parasitized erythrocytes per replicate) were centrifuged at room temperature (500 g), and the pellet was frozen at −80° C for downstream processing. Remaining cultures (3 technical replicates, each with or without Shld1) were supplemented with E64 (15 µM) for further culture until 55 hpi for collection as described above.

Further protein extraction steps for each of 10 samples were carried out in solutions supplemented with cOmplete protease inhibitors (Roche) and PhosSTOP phosphatase inhibitor cocktail (Roche). We released parasites with 0.05% saponin in PBS administered over several washes, for a total of ∼6.5 volumes of buffer for 1 packed erythrocyte volume of frozen pellet. Following additional washes in PBS without saponin, we added >1 volume of 8 M urea lysis buffer (100 mM NaCl, 25 mM Tris-HCl/pH 8), and subjected each sample to 5x freeze (−80°C)-thaw cycles before centrifugation at room temperature to separate protein-containing supernatant from pelleted cellular debris. Yields for each of the 10 samples ranged from 5-7 mg as assessed with the Pierce BCA (Bicinchoninic acid) protein assay. We reduced disulfide bonds with 5 mM tris(2-chloroethyl) phosphate (TCEP) for 30 min at room temperature, alkylated cysteines with 15 mM iodoacetamide for 30 min at room temperature in the dark, and quenched excess iodoacetamide by treatment with 5 mM dithiothreitol (DTT) for 15 min at room temperature. We precipitated protein in chloroform-methanol (*102*) before resuspension (8 M urea, 50 mM HEPES pH 8.5) and before dilution of urea to 1 M (50 mM HEPES pH 8.5) for digestion with LysC protease (1:100 protease-to-protein ratio, 3 h at 37 °C) before the addition of trypsin (1:100 protease-to-protein ratio) and continued digestion overnight at 37 °C. We quenched the reaction with 1% formic acid, carried out C18 solid-phase extraction (Sep-Pak, Waters), and precipitated peptides with vacuum-centrifugation.

#### Isobaric labeling with tandem mass tags (TMTs)

200 µg of peptides from each sample was dissolved in Buffer 1 (100 mM HEPES, pH 8.5). We carried out labelling with TMT reagents as previously described (*103*), according to manufacturer instructions (Thermo-Fisher Scientific). Following combination of the 10 TMT-labeled samples to match protein mass between samples, the mixture was vacuum-centrifuged and subjected to C18 solid-phase extraction (Sep-Pak, Waters) and eluate was collected.

#### Phosphopeptide enrichment

Peptides were resuspended in Buffer 1, followed by enrichment of phosphopeptides with “High-Select™ Fe-NTA Phosphopeptide Enrichment Kit” (Thermo-Scientific Cat. No. A32992) (*104*). The flow-through was retained for analysis of the proteome. Peptides and enriched phosphopeptides were dried by vacuum centrifugation.

#### Offline basic pH reversed-phase (BPRP) fraction

For proteomic analysis, the TMT-labeled peptide pool was fractionated via BPRP high-performance liquid chromatography (*103*). Eluted fractions were desalted, dried by vacuum-centrifugation, and resuspended in a solution of 5% acetonitrile and 5% formic acid, for mass spectrometry-based measurements.

For phosphopeptide analysis, we used the Pierce Off-line BPRP fractionation kit (Thermo Scientific), collecting and processing fractions for LC-MS/MS-based analysis as described previously (*13*).

#### Liquid chromatography and tandem mass spectrometry

We collected MS/MS data using an Orbitrap Fusion mass spectrometer (Thermo Fisher Scientific) coupled to a Proxeon EASY-nLC 1000 liquid chromatography (LC) pump (Thermo Fisher Scientific). For each analysis, 1 µg protein was loaded onto the LC onto an in-house pulled C18 column [30-35 cm, 2.6 um Accucore (Thermo Fisher), 100um ID] for MS-analysis (*13*).

Global proteome and phosphoproteome analyses each employed the multi-notch MS3-based method (*13, 105*). Global proteome and phosphoproteome analyses used an MS3-based TMT method (*106, 107*), which has been shown to reduce ion interference compared to MS2 quantification (*108*).

#### Mass spectrometry data analysis

Mass spectra were analyzed with a SEQUEST-based pipeline (*13, 109*). Peptide spectral matches (PSMs) were carried out a 1% false discovery rate (FDR), and filtered as previously described with minor modifications (*13*). To quantify the TMT-reporter ion, in each channel (0.003 Th range to distinguish reporters) we extracted the summed signal-to-noise (S/N) ratio and found the closest matching centroid to the expected mass. For proteomic analysis, PSMs (1% FDR) were collapsed to whole proteins (1% FDR). We used principles of parsimony to assemble the protein set, aiming to identify the smallest number of distinct polypeptides required to explain the observed PSMs. Relative protein levels were quantified by calculating the sum of reporter ion counts across associated PSMs (*109*). MS3 spectra represented in <2 TMT channels, and MS3 spectra with TMT reporter summed spectra of <100, or no MS3 spectra, were not considered for further analysis (*105*). Protein quantitation values were exported to Microsoft Excel. To normalize for variations in sample loading, within each TMT reporter ion channel, each parasite protein was normalized to the total signal of the *P. falciparum* proteome measured in that channel (*13*). For phosphoproteomics, the level of each phosphopeptide in a single TMT channel was normalized to the level of the parent protein in the same experimental conditions (i.e. average of normalized values for technical replicate measurements from proteomics). We did not apply this adjustment for the <2% of phosphopeptides that could not be mapped to a parent protein in proteomics.

#### Gene Ontology analysis of proteins upregulated with PfPP1-knockdown

For the 19 proteins shown to be increased in expression (>1.3-fold) in two proteomics measurements from separate experiments (Tables S1 and S2), we carried out gene ontology enrichment analysis using the webtool at PlasmoDB (http://www.plasmodb.org). The 10 most enriched GO categories are shown in Table S4.

#### Prioritization of late IDC-stage genes

For restriction of candidate substrates of *Pf*PP1 to proteins expressed late in the IDC, we used web-based tools at PlasmoDB (https://www.plasmodb.org) relying on published transcriptome data (*30*) to identify *P. falciparum* genes that are upregulated at the 40- or 48 hpi-mark of the IDC by at least 3-fold over levels at the midpoint of the IDC (average expression of 16 and 24 hpi), and by at least 1.5-fold over levels at 32 hpi.

### Chemical genetic assays

Compounds in DMSO (10 mM) were printed onto the surface of standard 96-well plates, using a D300e automated dispenser (Hewlett Packard), and stored at −20° C until addition of parasites. Chemicals in water were added individually onto the surface of 96-well plates, or added directly at the high concentration to parasites in wells before serial dilution in-plate. Phospholipids (or methanol vehicle) at 10x final concentration were added to the surface of plates and allowed to evaporate before addition of parasites. Synchronized *Pf*PP1-DD parasites were washed extensively in media without Shld1 and supplemented with varying concentrations of Shld1 (constant volume of ethanol carrier across doses), as indicated in the data. Parasites with Shld1, set at 0.5%-hematocrit, were added at 100 µl volume to wells with compounds to attain the reported final concentrations. Wells immediately surrounding the samples were filled with water or aqueous solution to prevent evaporation through the course of the assay. Parasites were allowed to incubate with inhibitor for ∼20 hrs before fixation and measurement of egress from schizonts and reinvasion to rings by flow-cytometry.

## Data availability

All data are available in the manuscript or supplementary materials.

## Acknowledgements

We thank the Harvard Electron Microscopy Facility and the Montpellier Ressources Imagerie for imaging. We thank the laboratories of Drs. Jeffrey Dvorin, Dyann Wirth, Barbara Burleigh, for use of equipment and reagents, as well as discussions. We thank Drs. Marcus Lee, Anthony Holder and Michael Blackman for strains and reagents. This work was supported by NIH R21 AI128480 (M.T.D.), NIH R01 AI138551 (M.T.D.), NIH GM132129 (J.A.P.), NIH GM67945 (S.P.G), Laboratoire d’Excellence (LabEx) EpiGenMed ANR-10-LABX-12-01 (M.L., C.B.B.), Fondation pour la Recherche Médicale (Equipe FRM EQ20170336725, M.L.).

## Author Contributions

A.S.P., A.M., J.A.P., A.M.B., L.B., M.S., A.L.K., B.E., J.S.N.A, and M.H.L, carried out experiments; A.S.P., A.M., J.M.G., M.S., C.B.B., and M.H.L., analyzed the data; A.S.P., J.A.P., S.P.G., M.H.L., and M.T.D., designed experiments; A.S.P., M.H.L., and M.T.D., wrote the manuscript.

## Supplementary Information

### Supplementary Text

#### Assessment of the specificity of PfPP1-function according to phosphoproteomic data

At 48-hpi, 30 phosphopeptides (0.8% of 3884 total detected) exhibit a reduction of >2-fold with *Pf*PP1-DD destabilization, and 60 (1.5% of total) are increased by >2-fold (Table S3). Other evidence suggests limited effects secondary to perturbation of *Pf*PP1-function at 48-hpi: (i) proliferation defects elicited by knockdown are reversible to a large degree with resupplementation of Shld1 at 48-hpi, but not 55-hpi (Fig. S4A); (ii) in contrast to 55-hpi, changes in the global phosphoproteome with *Pf*PP1-DD knockdown at 48-hpi are not predictive of further developmental progression (Fig. 3D; fig. S4B); and (iii) overall variability in the phosphopeptide levels at 48-hpi, but not 55-hpi, induced by *Pf*PP1-DD-destabilization is comparable to a baseline, determined from the distribution of non-phosphorylated peptides in the same sample (Fig. S4C) (*110*). Of the phosphopeptides upregulated by >2-fold with *Pf*PP1-DD knockdown at 48-hpi (y-axis, Fig. 3D), we regarded sites that did not decrease with IDC progression from 48 to 55-hpi (x-axis, Fig. 3D) to be least likely to be influenced by effects secondary to perturbation of *Pf*PP1.

**Figure S1.**
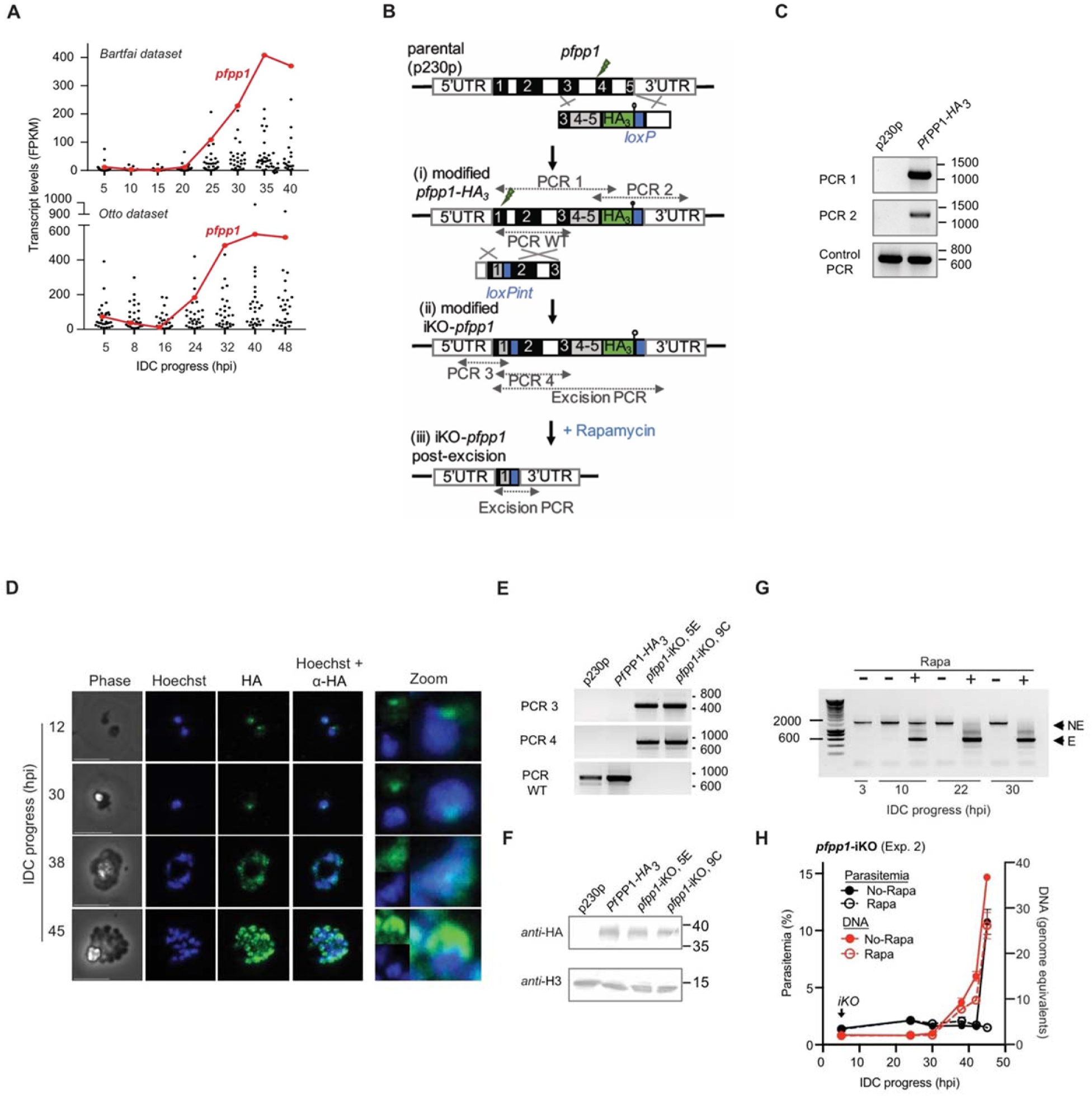
**(A)** For 29 *P. falciparum* protein phosphatase genes (*8*), levels of mRNA expression (frequency per kilobase per millions reads, fpkm) over the course of the IDC measured in two separate transcriptome-wide studies from Bártfai and colleagues (top), and Otto and colleagues (bottom) (*29, 30*). **(B)** Sequential double-crossover recombination at the *pfpp1* chromosomal locus in the DiCre recombinase-expressing p230p parental parasite line to generate (i) the *Pf*PP1-HA_3_ line with a 3’ loxP site; and (ii) the *pfpp1*-iKO with a loxP-containing synthetic intron (loxPint); followed by (iii) treatment with rapamycin to induce excision of *pfpp1*. Sites for PCR amplicons to confirm construction of transgenic lines are indicated. Black and white bars represent exons and introns of the *pfpp1* gene, respectively. Grey bars represent recodonized region of the *pfpp1* gene and blue bars stand for loxP site. Lollipop: stop codon; green lightning bolt: Cas9 double strand break. **(C)** PCRs to confirm construction of *Pf*PP1-HA_3_ with 3’-loxP, as in (B). **(D)** Localization of *Pf*PP1-HA_3_, relative to Hoechst-stained nuclei, assessed by immunofluorescence. Staging is indicated by IDC progress. **(E)** PCRs to confirm construction of *pfpp1*-iKO with loxP-containing synthetic intron, as in (B). PCR also confirm unmodified *pfpp1* (PCR WT). **(F)** Immunoblot performed with anti-HA antibodies to show that *Pf*PP1-HA_3_-loxP and *pfpp1*-iKO parasite lines express a tagged version of the protein of the same molecular mass. Anti-histone H3 was used as a loading control. **(G)** PCRs to confirm Rapa-induced excision of the *pfpp1* gene in *pfpp1*-iKO transgenic line following rapamycin treatment at 3 hpi. NE and E indicate the non excised (1606 bp) and excised (581 bp) versions of *pfpp1* gene, respectively. Sizes are indicated on the left in bp. **(H)** A second experiment to measure the effect on DNA replication and proliferation in *pfpp1-iKO* parasites following +/− Rapa-treatment at 5 hpi, as in Fig. 1C.

**Figure S2.**
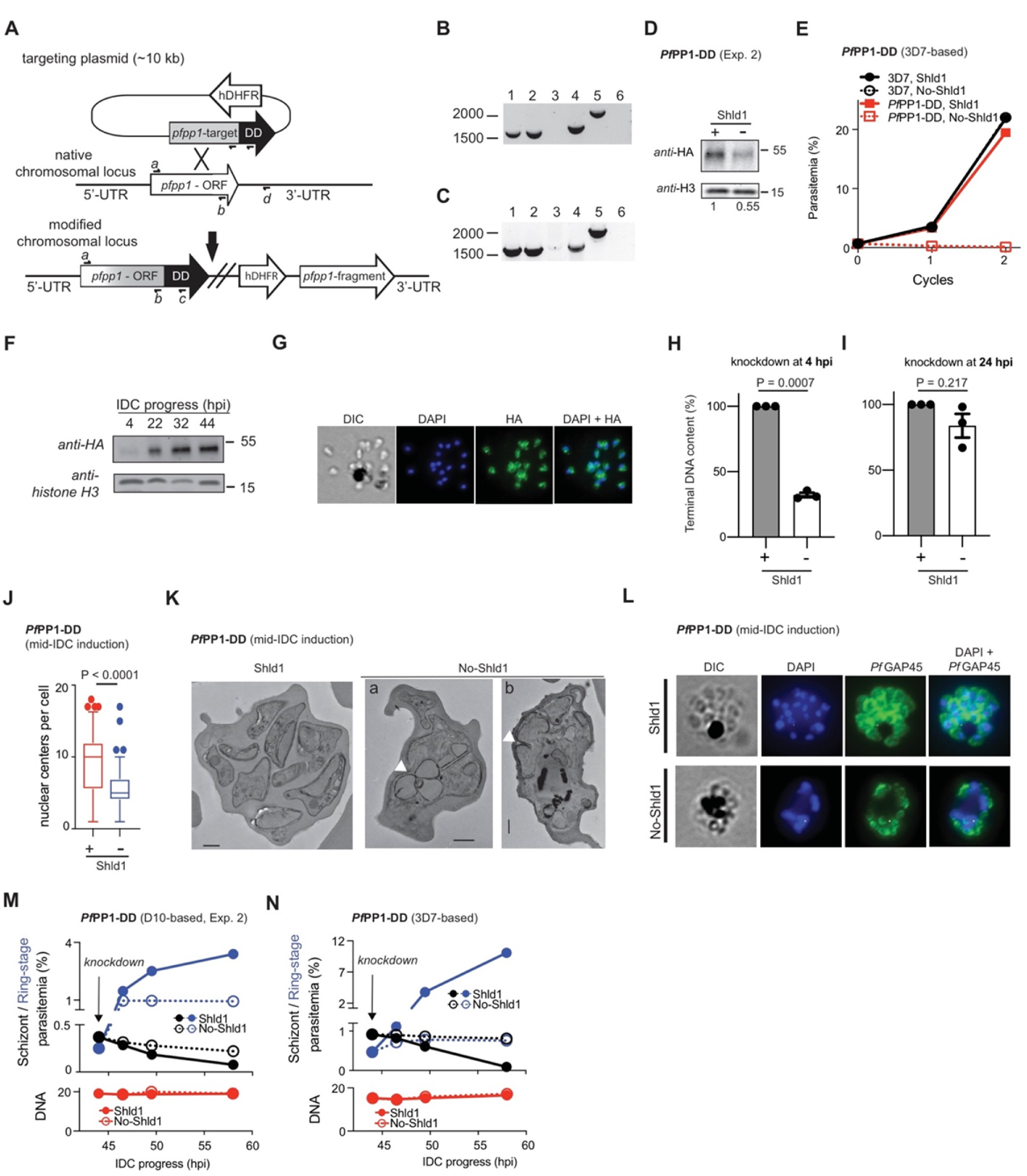
**(A)** Top: Scheme for single-crossover integration of the *dd-tag* into the endogenous *pfpp1* genetic locus, with binding sites for diagnostic primers indicated. **(B-C)** PCR to confirm integration of *dd* into *pfpp1* in the D10 (B) and 3D7 background strains (C). In electrophoresis gel, lanes 1,3,5: wild type parasite background; lanes 2,4,6: clonal *Pf*PP1-DD transgenic parasites. Lanes 1,2: diagnostic primers *a,b*; lanes 3,4: primers *a,c*; lanes 5,6: primers *a,d*. Positions by size (bp) are indicated. **(D)** An immunoblot as in Fig. 1D, schizont-stage *Pf*PP1-DD parasites grown +/-0.2 µM Shld1 for ∼20 hrs. **(E)** Proliferation of *Pf*PP1-DD in the 3D7 background, assessed as in Fig. 1E. **(F)** *Pf*PP1-DD expression in synchronized parasites in 0.2 µM Shld1 at the indicated timepoints following erythrocyte invasion, assessed with anti-HA tag antibody. Histone H3 levels were measured for loading control. **(G)** Assessment of *Pf*PP1 localization in a recently ruptured *Pf*PP1-DD schizont-stage parasite using an antibody for HA. Nuclear staining was carried out with DAPI. **(H-I)** DNA content in advanced-stage parasites (51-55 hpi) +/− induction of *Pf*PP1-DD knockdown early (4 hpi, H) or at a mid-stage of the IDC (24 hpi, I), assessed by flow-cytometry. **(J)** Box plot of numbers of nuclear centers (median, outliers at >95% percentile with none observed below 5%) in terminally developed parasites with induction of *Pf*PP1-DD knockdown at a mid-stage of the IDC (experiment from Fig. 1F, middle panel). Two-tailed Mann-Whitney test. **(K)** Electron microscopy images of terminally developed *Pf*PP1-DD parasites (55 hpi), with knockdown induced at 24 hpi (+/-0.3 µM Shld1). Parasites were supplemented with 50 µM E64 at ∼47 hpi to block egress. For knockdown-cells (No-Shld1), we show parasites arrested at a late stage when incipient cytokinesis in the vicinity of dividing nuclei are apparent (white arrowheads). **(L)** Immunofluorescence microscopy images of terminally developed *Pf*PP1-DD parasite cells, with knockdown and sampling as in (K), indicating the inner membrane complex marker *Pf*GAP45. **(M-N)** Egress and erythrocyte re-invasion following knockdown of *Pf*PP1-DD in either the D10 (M) or the 3D7 background (N) at 44 hpi, and DNA content in E64-treated parasites (50 µM) as described in Fig. 1H.

**Figure S3.**
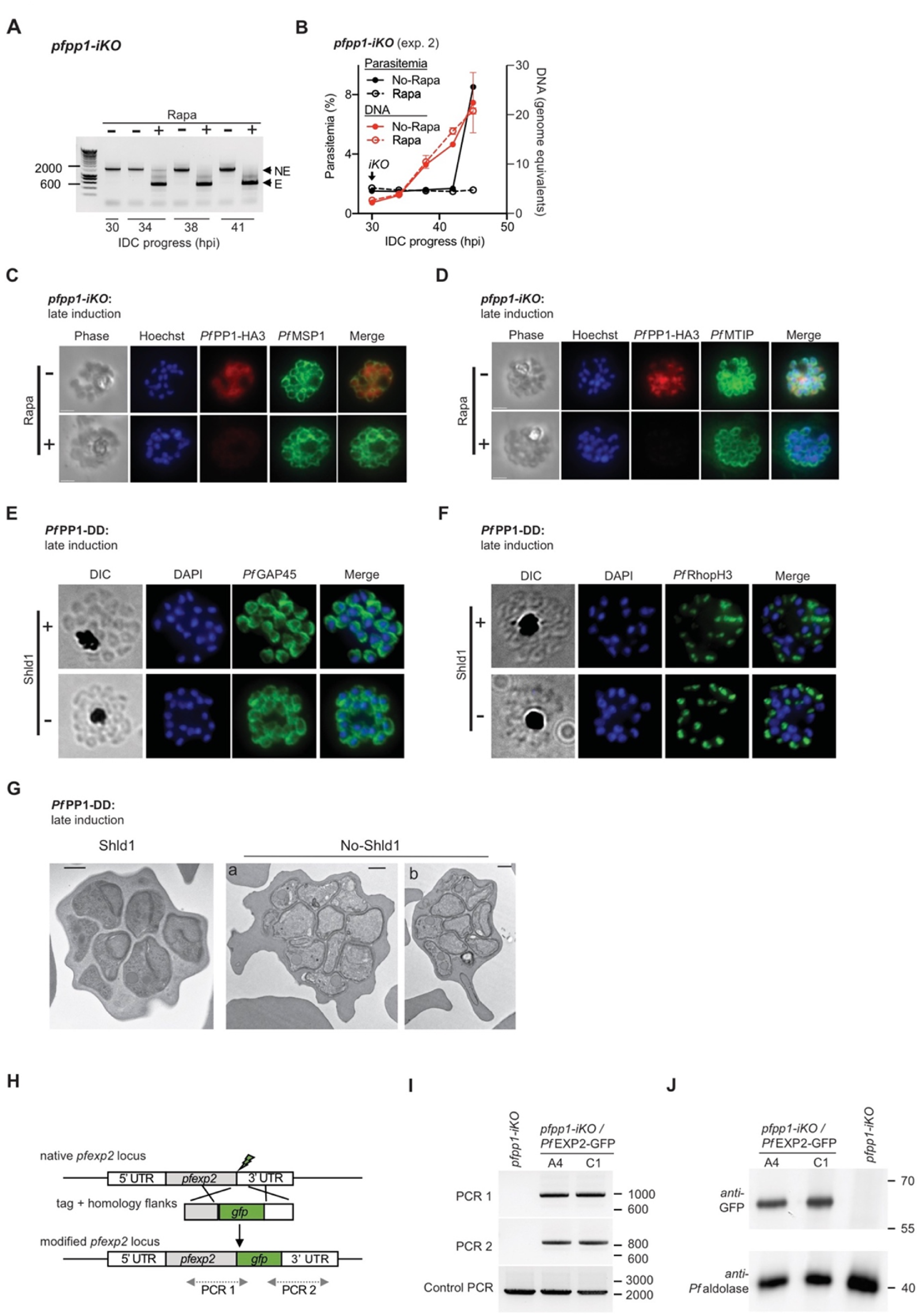
**(A)** PCR to confirm excision of the *pfpp1* gene following iKO with rapamycin at 30hpi (trophozoite stage), as in fig. S1G. **(B)** A second experiment measuring parasitemia and DNA synthesis following iKO of *pfpp1*, as in Fig. 2B. **(C-D)** Immunofluorescence images showing depletion of *Pf*PP1-HA_3_ in parasites assessed in Figs. 2E (C) and 2F (D). **(E-F)** Immunofluorescence microscopy images of *Pf*PP1-DD parasite cells, sampled at 55 hpi, indicating the inner membrane complex marker *Pf*GAP45 (E) or the rhoptry marker *Pf*RhopH3 (F), with knockdown (+/− 0.5 µM Shld1) induced at 44 hpi. Parasites were supplemented with 50 µM E64 to block egress. **(G)** Electron microscopy images of terminally developed *Pf*PP1-DD parasites treated as in (E-F). **(H)** Modification of the native *pfexp2* locus for 3’-tagging with GFP in the *pfpp1-*iKO background, via Cas9-mediated double-stranded break repair (see fig. S1B). **(I)** PCRs to confirm construction of *pfpp1*-iKO */ Pf*EXP2-GFP, as in (H). **(J)** Immunoblot to show GFP expression in *pfpp1-iKO / Pf*EXP2-GFP lines. *Pf*aldoase was assessed to control for loading.

**Figure S4.**
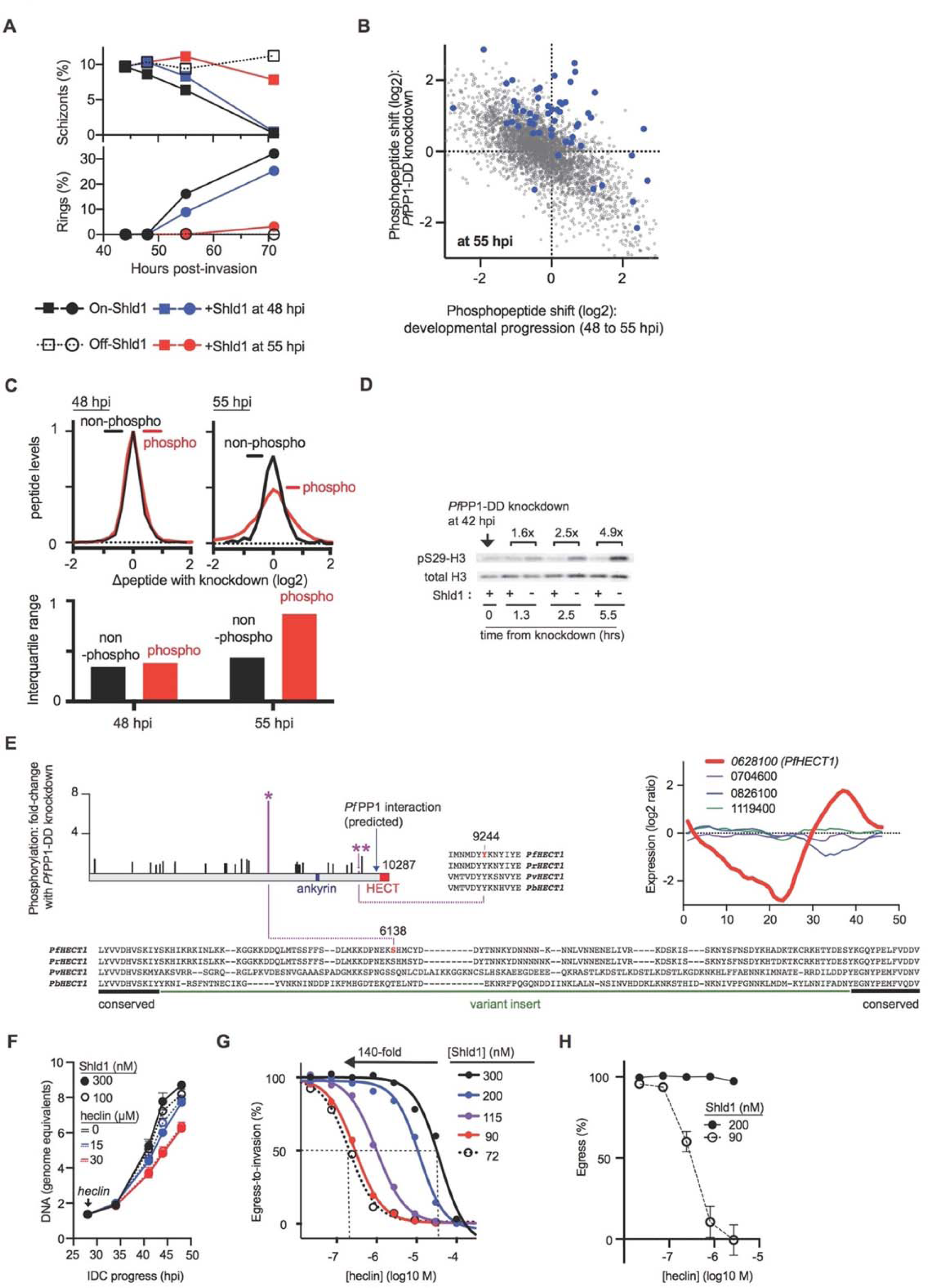
**(A)** Timecourse of schizont and ring-stage parasitemia in control samples for phosphoproteomic analysis of *Pf*PP1-DD knockdown, assessed from thin blood smears. Parasites were maintained continuously on-Shld1 (0.3 µM) or off-Shld1 (EtOH) following start of experiment at 44 hpi. To assess reversibility of the knockdown-phenotype, off-Shld1 parasites were supplemented with Shld1 (1 µM) at the two timepoints for sample collection. **(B)** The effect of *Pf*PP1-DD-based perturbation on levels of all detected phosphopeptides at 55-hpi plotted against changes with development (+Shld1-parasites) from 48 to 55-hpi, as in Fig. 3D. All phosphopeptides upregulated by >2-fold at 48-hpi by *Pf*PP1-DD knockdown (Fig. 3D) are indicated in blue, showing that these hits are masked by developmental effects dominant by 55-hpi. **(C)** Top: Following TiO2-based affinity purification, the distribution of the log2-fold differences with *Pf*PP1-DD knockdown for either the enriched phosphopeptides (red) or non-phosphorylated peptides (black) measured by mass-spectrometry. Data for samples collected at both the 48 (left) and 55-hpi timepoints (right) are shown. Bottom: The interquartile ranges for data shown in top panel. **(D)** Phospho-histone H3 (Ser-29) dynamics with *Pf*PP1-knockdown in parasites late in the IDC, assessed by immunoblot analysis. The timepoint of *Pf*PP1-knockdown and timepoints for sampling following knockdown are indicated. The fold-increase in phospho-histone H3 (S29), normalized to total histone H3, is indicated above the blot (mean of 2 technical replicate timecourses). **(E)** Left: Schematic of *Pf*HECT1 as in Fig. 3E with *Pf*PP1-regulated phosphosites shown in sequence alignment with orthologs from other *Plasmodium* spp. (*P. reichenowi*, *Pr; P. vivax*, *Pv; P. berghei, Pb*). For Ser-6138, extended sequence is shown to highlight regions of conservation and variation. Position of the predicted interaction site for *Pf*PP1 (*34*) is shown (blue arrow). Right: Relative transcript expression over the course of IDC (*111*) for the 4 HECT domain-containing proteins in *P. falciparum*, each denoted by the 7-digit suffix of the systematic gene ID beginning with “PF3D7_”. **(F)** DNA replication in *Pf*PP1-DD parasites following administration of heclin at 28-hpi with indicated concentrations (DMSO control, black, 15 µM, blue; 30 µM, red;), and growth supported by a high (300 nM) or suboptimal (100 nM) concentration of Shld1 (mean +/− s.e.m.; n=3 experiments). **(G)** Dose-response curves from a single experiment to measure susceptibility of egress-to-invasion by *Pf*PP1-DD parasites to heclin at various Shld1 concentrations, with range of IC50 values indicated. **(H)** The susceptibility of egress by *Pf*PP1-DD parasites to heclin at 200 or 90 nM Shld1 (mean +/− s.e.m.; n=3 experiments).

**Figure S5.**
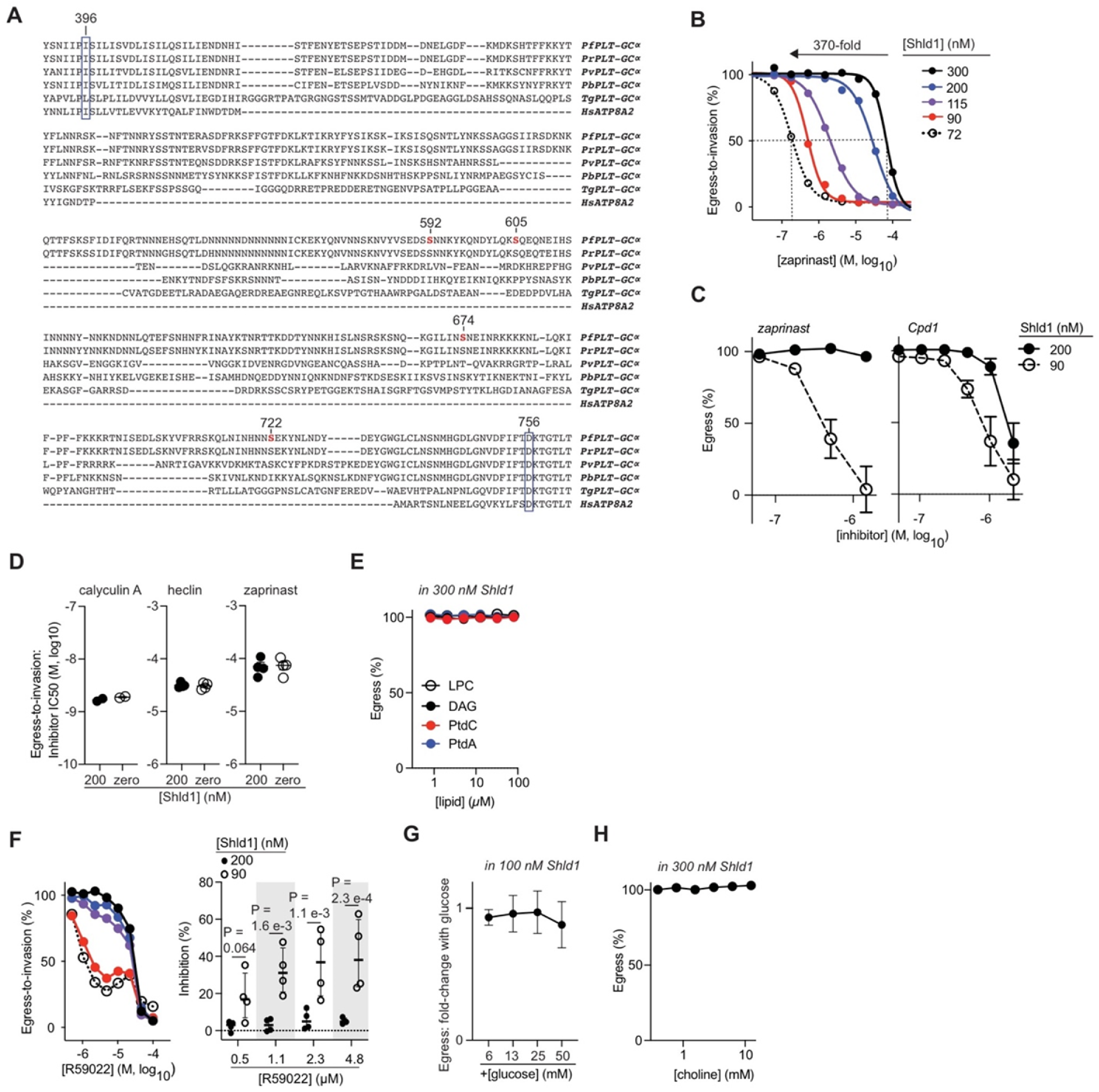
**(A)** Alignment of the multi-phosphorylated cytoplasmic domain between the fourth and fifth transmembrane domains of the PLT in *Pf*PLT-GCα (see Fig. 4A). The alignment shows orthologs from *P. reichenowi* (*Pr*), *P. vivax* (*Pv*), *P. berghei* (*Pb*), and *T. gondii* (*Tg*); and also the human protein ATP8A2 (UniProt Q9NTI2.2) to show conservation of a isoleucine residue (396 in *Pf*PLT-GCα) in transmembrane 4 required for transport of phospholipids (*112*), indicated in purple. *Pf*PP1-responsive phosphorylation sites are indicated in red. The absolutely conserved catalytic aspartate residue in the consensus sequence DKTGTLT required for ATPase-dependent phospholipid transport is indicated in purple. **(B)** An experiment with dose-responses of *Pf*PP1-DD parasites to zaprinast (egress-to-invasion) elicited over a multiple Shld1 concentrations, with range of IC50 values indicated. **(C)** Sensitivity of egress by *Pf*PP1-DD parasites to zaprinast (left) or Cpd1 (right). **(D)** The sensitivity of egress-to-invasion by D10 parasites (parent to *Pf*PP1-DD) for the indicated compounds at the indicated Shld1 concentrations (mean +/− s.e.m.; n=2-4 experiments as indicated). **(E)** In 300 nM Shld1, percent egress for *Pf*PP1-DD parasites in the presence of the lipids in Fig. 4B at the indicated concentrations (mean +/− s.e.m.; n=4 experiments). **(F)** Left: A series of dose-response curves in a single experiment showing induced sensitivity of parasite egress-to-invasion to R59022 with knockdown of *Pf*PP1-DD at concentrations <10 µM. Shld1 concentrations as in (B). Right: Induced sensitivity of parasites to low concentrations (<10 µM) of R59022 with partial knockdown of *Pf*PP1-DD, based on inhibition of egress-to-invasion (mean +/− s.e.m.; n=4 experiments). Significance assessed by multiple *t-*tests. **(G)** In 100 nM Shld1, the effect of supplementation with the indicated concentrations of glucose on egress by *Pf*PP1-DD parasites, expressed as fold-change compared to no-added glucose (mean +/− s.e.m.; n=4 experiments). The baseline concentration of glucose in media is 11.1 mM. **(H)** In 300 nM Shld1, percent egress for choline (see Fig. 4E) at indicated concentrations, as in (E) (mean +/− s.e.m.; n=4 experiments).

**Figure S6.**
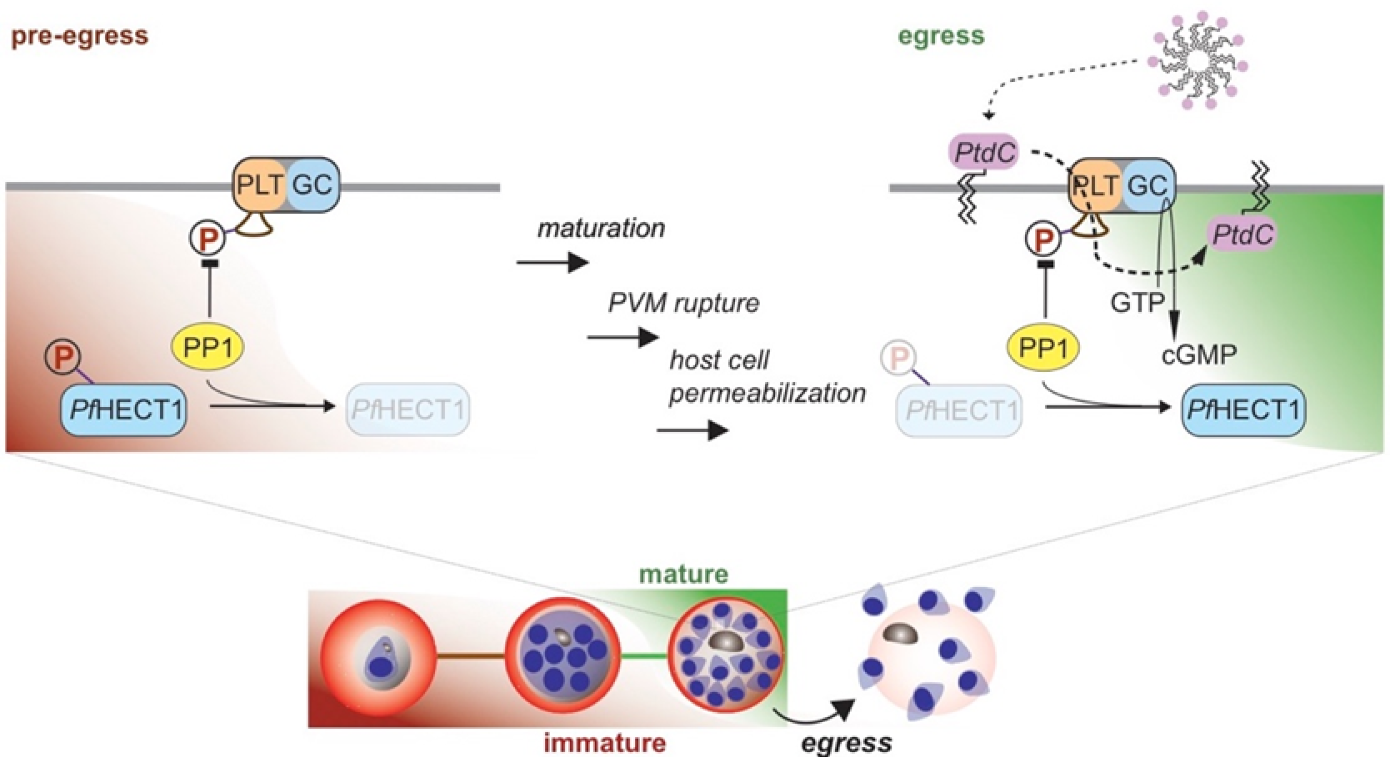
*Pf*PP1-regulated cellular pathways in coordination with late IDC-stage developmental progression of *P. falciparum*. At an immature state when the parasite is not yet competent to process signals for egress (pre-egress), *Pf*PP1 acts on *Pf*PLT-GCα to suppress cGMP levels. *Pf*PP1 acts on *Pf*HECT1 as the parasite matures, and cGMP activates the egress program leading to PVM rupture and host cell permeabilization. We propose that PtdC from the serum is translocated by *Pf*PLT-GCα, providing a late signal to stimulate egress.

**Table S1**. Proteomic analysis of *Pf*PP1-DD parasites in late IDC-stage parasites.

**Table S2**. A second proteomic analysis of *Pf*PP1-DD parasites in late IDC-stage parasites.

**Table S3**. Phosphoproteomic analysis of *Pf*PP1-DD parasites in late IDC-stage parasites.

**Table S4**. Gene Ontology analysis of proteins upregulated with *Pf*PP1-DD knockdown in late IDC-stage parasites.

**Table S5**. Primers used in this study.

